# Recreation of an antigen-driven germinal center in vitro by providing B cells with phagocytic antigen

**DOI:** 10.1101/2020.02.26.965921

**Authors:** Ana Martínez-Riaño, Pilar Delgado, Pilar Mendoza, Clara L. Oeste, David Abia, Elena Rodríguez-Bovolenta, Martin Turner, Balbino Alarcón

## Abstract

Successful vaccines rely on activating a functional humoral response that results from generating class-switched high affinity immunoglobulins (Igs) with a superior capacity to neutralize infection. Key to this process is the germinal center (GC) reaction, in which B cells are selected in their search for antigen and T cell help. A major hurdle to understanding the mechanisms of B cell:T cell cooperation has been the lack of an *in vitro* system to recreate GCs in an antigen-specific manner. Here we report the generation of functional antigen-specific high affinity Igs of different isotypes in simple 2-cell type cultures of naïve B and T cells. It is crucial for this process for B cells to take up antigen by a phagocytic mechanism, which results in stronger and more sustained BCR signals compared to stimulation with a soluble antigen. We also show the applicability of the system to generate antibodies of potential clinical interest.

## INTRODUCTION

For an efficient protective humoral response to pathogen-derived protein antigens, B cells establish an intimate collaboration with antigen-specific helper T cells. To obtain T cell help, B cells have to recognize cognate antigen via their B cell antigen receptor (BCR), internalize it and present it once processed as MHC class II associated peptides. CD4 T cells that are able to recognize the processed antigen will become activated and express ligands of costimulatory receptors for B cells that, in turn, will initiate immunoglobulin class-switching, proliferation and somatic hypermutation. These processes result in the selection of B cells bearing class-switched immunoglobulins of high affinity for the antigen. This B-T cell cooperation takes place in germinal centers (GC), where B cells undergo iterative cycles of antigen recognition and presentation to T cells followed by very rapid cell proliferation and expansion. It is generally accepted that in GCs, B cells establish a fierce competition for antigen to gain T cell help resulting in the selection of B cells bearing BCRs with the highest affinity (Gitlin et al., 2014). The BCR can interact and be activated by soluble proteins although it is believed that, most frequently, B cells recognize and take up antigen deposited on the surface of antigen-presenting follicular dendritic cells (Avalos and Ploegh, 2014; Phan et al., 2009; Suzuki et al., 2009). It has long been thought that only antigen-presenting cells of myeloid origin are able to phagocytose antigens and that B cells are not competent to phagocytose particulate antigens (Ochando et al., 2006; Vidard et al., 1996). However, growing evidence suggests that B cells can also perform phagocytic functions. Antigen phagocytosis by B cells was first described in early vertebrates (Li et al., 2006; Zimmerman et al., 2010), but lately it has also been demonstrated that murine B-1 B cell populations and human liver B cells can phagocytose bacteria (Gao et al., 2012; Nakashima et al., 2012; Parra et al., 2012; Zhu et al., 2016).

B cells receive help from a type of activated helper CD4 T cell known as T follicular helper cells (TFH). These cells release important cytokines that stimulate B cell proliferation and modulate Ig class switching, including IL4 and IL21. Furthermore, TFH express ligands (CD40L, ICOS) for costimulatory receptors in B cells (CD40 and ICOSL) (Ramiscal and Vinuesa, 2013). B cells integrate signals emanating from their antigen-engaged BCR, from ligated CD40 and ICOSL, as well as from cytokine receptors to promote their program of affinity maturation and Ig class switching. In this context, the BCR has a dual function, first as a provider of activation signals to the B cell and second as a mediator of antigen internalization, processing and presentation to T cells. It is a challenge to distinguish the extent to which BCR function depends indirectly on its antigen presentation role, and thereby on the signals transmitted by CD40 and other T cell-engaged costimulatory receptors, from the contribution of BCR signaling (Nowosad et al., 2016). One of the caveats in studying the molecular processes that take place during B-T cell interaction in detail is the inability to recreate GC *in vitro*. Different protocols consisting of mixtures of cytokines and the expression of CD40L in non-T cells have been used (Nojima et al., 2011). However, these procedures are not antigen-specific and have not allowed the selection for Ig class-switched B cells with increasing affinity for antigen.

Receptor-mediated phagocytosis of particulate material requires an actin-dependent zippering of membrane around the particle, forming a cup that leads to progressive engulfment (Groves et al., 2008). Phagocytosis is regulated by GTPases of the Rho family. According to the involvement of different Rho family members, phagocytosis is classified into two main groups. Type I phagocytosis involves Rac1 and Cdc42, such as that observed for the Fc Receptor (FcR). In turn, type II phagocytosis involves RhoA, as described for the Complement Receptor 3 (CR3; reviewed in (Niedergang et al., 2016). Another important GTPase is RhoG, which is an evolutionarily conserved, intracellular mediator of apoptotic cell phagocytosis (deBakker et al., 2004; Henson, 2005). Interestingly, in an RNA interference screen of 20 Rho GTPases in macrophages, RhoG was found to be required for particle uptake mediated by both FcR and C3R (Tzircotis et al., 2011). At odds with the idea that lymphocytes are not professional phagocytes, we previously found that RhoG was involved in the nibbling of MHC-associated portions of the membrane of antigen-presenting cells by T cells (Martinez-Martin et al., 2011). This process, known as trogocytosis, also requires the activation of another small GTPase, in this case of the R-Ras subfamily, known as R-Ras2 or TC21, which is a direct interactor of the T cell receptor (TCR). In addition, we found that T cells can phagocytose 1-6 μm diameter latex beads coated with anti-CD3 antibodies by a RRas2 and RhoG-mediated process (Martinez-Martin et al., 2011). The capacity of T cells to phagocytose particles by a TCR-driven mechanism is paralleled by a similar behavior of B cells. We have recently shown that naive follicular B cells can phagocytose 1-3 μm beads coated with their cognate antigen and that such process is BCR-driven and leads to the presentation of antigen to T cells which subsequently engage in several rounds of cell divisions (Martinez-Riano et al., 2018). Furthermore, such process of BCR-driven antigen phagocytosis leads to the efficient generation of germinal centers (GC) *in vivo* as well as to the generation of antigen-specific class-switched Igs of high affinity. Having demonstrated the stimulatory effect of antigen phagocytosis by B cells on GC formation in vivo, we have now investigated its potential use for the recreation of antigen-driven GCs in vitro.

## RESULTS

### Generation of antigen-specific GC B cells in vitro by a phagocytic-dependent mechanism

Here, we investigated the possibility of using antigen phagocytosis by B cells to recreate a GC *in vitro*. We first tested if naïve follicular B cells expressed markers of GC B cells when stimulated with a polyclonal BCR stimulus in the presence of helper CD4+ T cells. We found that WT B cells incubated for 3 days with 1 μm beads coated with anti-IgM and ovalbumin, in the presence of ovalbumin-specific OT2 TCR transgenic T cells, proliferated and expressed the GC B cell markers CD95 and GL7 (Fig. 1A). They also upregulated CD40 and their proliferation depended on T cell help and RhoG expression (Fig. 1A, 1B and Suppl. Fig. 1A). In parallel, we evaluated the effect of antigen phagocytosis by B cells on T cell activation and differentiation. Preincubation of B cells with beads coated with anti-IgM and OVA induced the upregulation of IL-2Rα (CD25) expression by OT-2 T cells, whereas preincubation with beads coated with either anti-IgM or OVA alone did not (Fig. 1C). These data suggest that B cell activation alone is not sufficient to activate CD4 T cells, but rather that OVA antigen uptake by B cells requires a BCR-dependent process. Furthermore, compared to WT B cells, CD25 expression by OT-2 T cells was reduced if B cells lacked RhoG (Fig. 1C). In addition, we found that OT-2 T cells expressed CXCR5 and PD1, two markers of CD4 T cell differentiation towards follicular helper T cells (TFH) (Ramiscal and Vinuesa, 2013), when stimulated with B cells that had been pre-incubated with 1 μm beads coated with anti-IgM and OVA but not with anti-IgM or OVA alone (Fig. 1D). Again, expression of the TFH markers by OT2 was reduced if B cells lacked RhoG and were therefore deficient in antigen phagocytosis. These data suggested that antibody-mediated BCR triggering can promote antigen phagocytosis and presentation to T cells, inducing T cell and B cell proliferation as well as the acquisition of B cell and T cell markers typical of a GC response.

**Figure 1.**
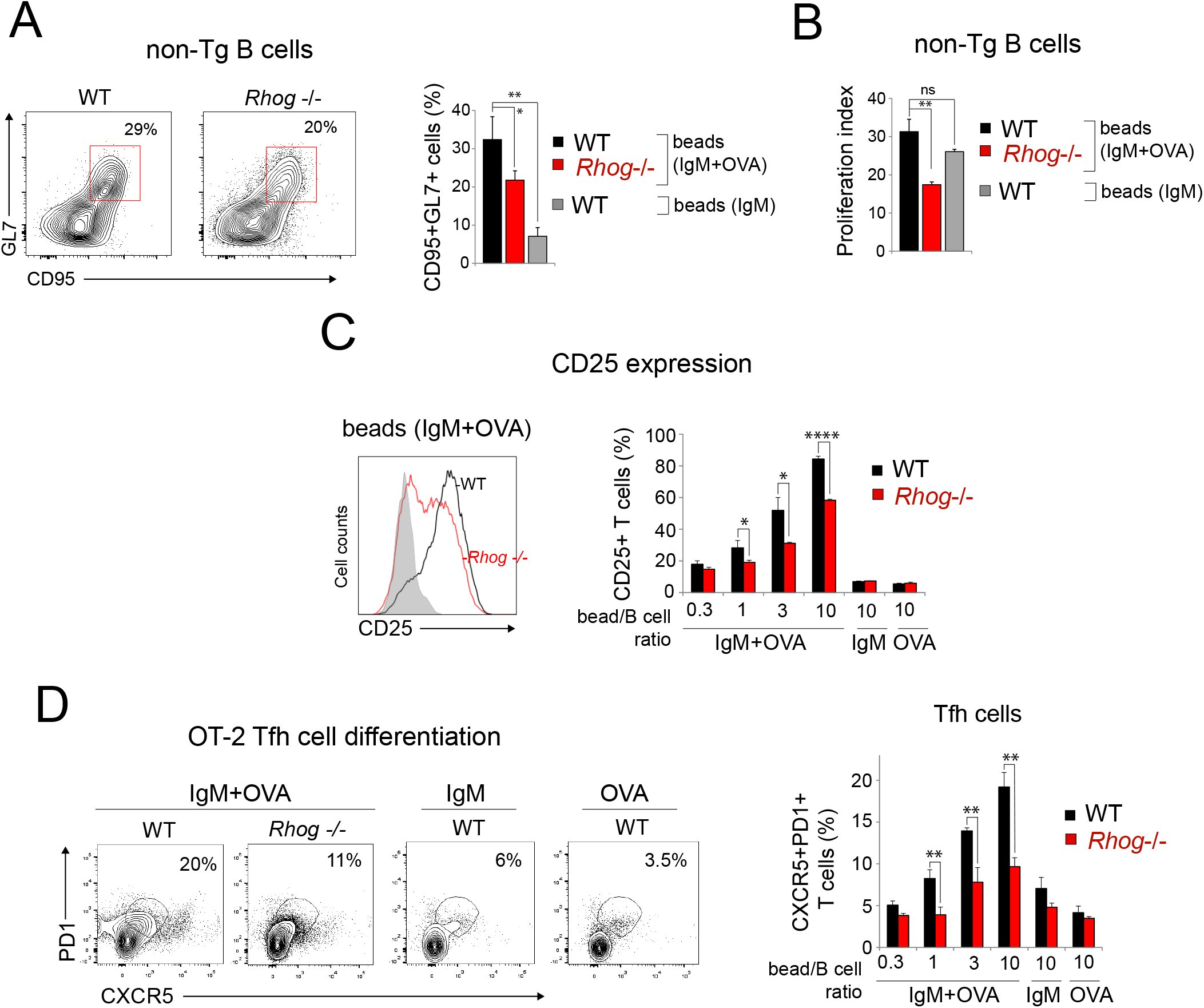
B cells and T cells differentiate in vitro into GC B cells and TFH upon B cell activation with antibody-coated beads. **(A)** Naïve B cells from WT and *Rhog*^−/−^ mice were preincubated with 1 μm beads coated with IgM plus OVA or IgM alone, at a 20:1 bead/cell ratio, and co-cultured for 3 days with OT-2 T cells (1:1 B/T cell ratio). FACS contour plots to the left show the appearance of a double positive (CD95^+^ GL7^+^) population in gated B220^+^ B cells. The bar plot to the right represents mean ± S.D. (*n* = 3). * *p*<0.05; ** *p*<0.005 (unpaired Student’s t test). **(B)** Proliferation of B cells after 3 days of culture was calculated by CTV dilution. Data represent the mean ± S.D. (*n* = 3). ** *p*<0.05; ** *p*<0.005 (unpaired Student’s t test). **(C)** Induction of CD25 expression by OT-2 T cells incubated with phagocytic B cells as in (A) for 3 days. The histogram to the left shows an overlay of CD25 expression in OT-2 cells incubated with: wild type B cells preincubated with beads coated with anti-IgM plus OVA (black line), with RhoG-deficient B cells preincubated with anti-IgM plus OVA (red line), or with wild type B cells preincubated with beads coated with anti-IgM alone (grey shaded). Bar plot to the right represents mean ± S.D. (*n* = 3). * *p*<0.05; ** *p*<0.005; **** *p*<0.00005 (unpaired Student’s t test). **(D)** Induction of TFH marker (PD1 and CXCR5) expression in OT-2 T cells after 3 days of culture with WT or RhoG-deficient B cells as in (C). Bar plots represent the percentage of double positive OT-2 T cells. Data represent the mean ± S.D. (*n* = 3). ** *p*<0.005 (unpaired Student’s t test).

To determine if GC B cell differentiation was induced upon phagocytosis in an antigen-specific manner, we co-cultured B cells isolated from B1-8^hi^ BCR knock-in mice with OT-2 T cells. B1-8^hi^ knock-in mice bear a rearranged VDJ region in the IgH locus that, in combination with a rearranging lambda light chain, confers specificity for the hapten 4-hydroxy-3-nitrophenylacetyl (NP) and its iodinated derivative 4-hydroxy-3-iodo-5-nitrophenylacetic acid (NIP). We incubated purified B1-8^hi^ B cells with 1 μm beads coated with NIP-OVA in the presence of OT-2 T cells for 4 days. This led to the emergence of GC B cells characterized by the expression of the GL7 and CD95 markers and to B cell proliferation (Fig. 2A and 2B). Beads coated with NP linked to a different carrier protein (chicken gammaglobulin, CGG) did not elicit GC B cell differentiation or proliferation, indicating that T cell help is required. Likewise, the acquisition of a GC B cell phenotype was inhibited if B cells lacked RhoG, suggesting that beads, and the NIP-OVA antigen, were taken up by phagocytosis.

**Figure 2.**
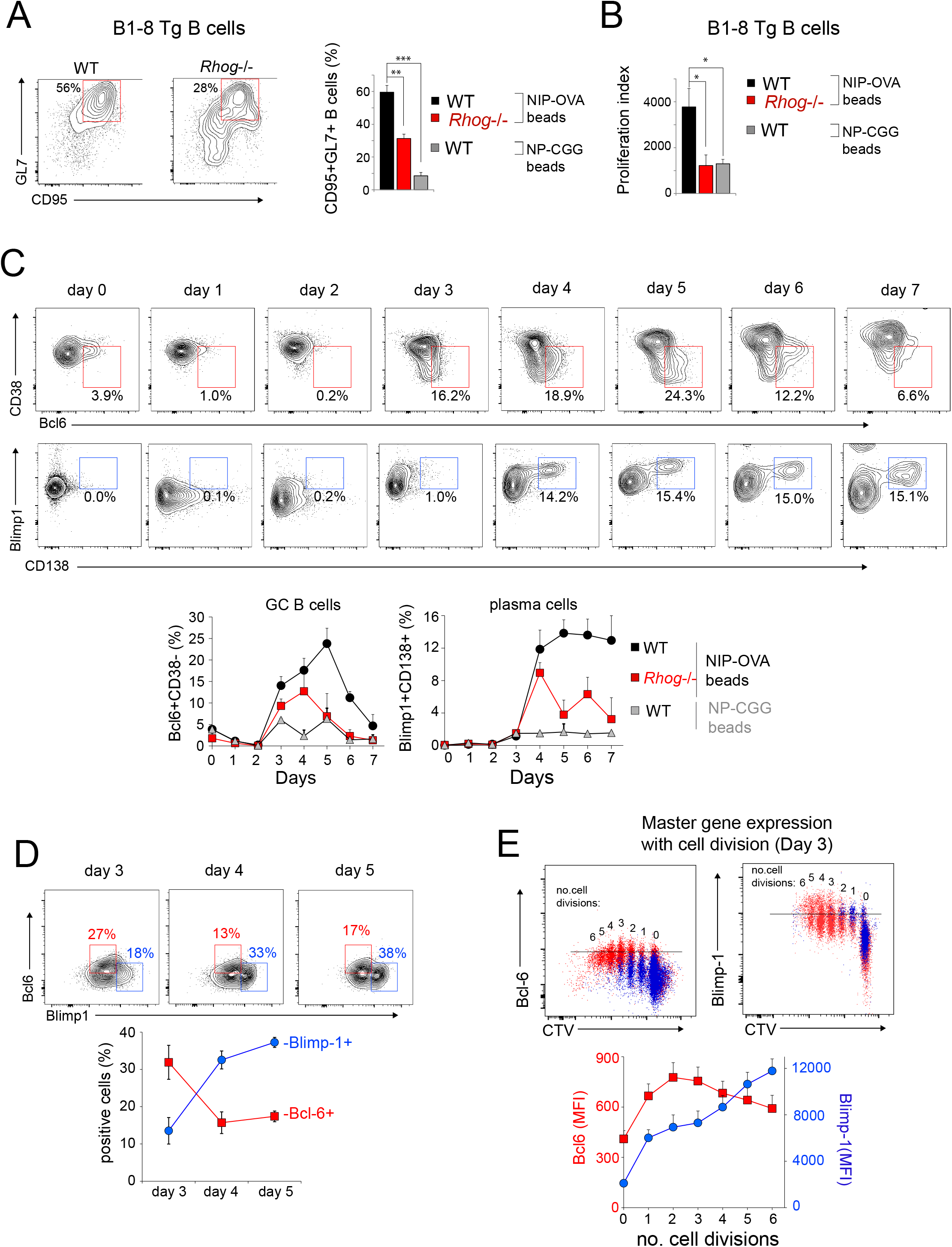
Phagocytic B cells sequentially differentiate in vitro into GC B cells and plasma cells upon B cell activation with antibody-coated beads. **(A)** Naïve B cells from B1-8^hi^ transgenic WT and *Rhog*^−/−^ mice were preincubated with 1 μm beads coated with NIP-OVA or NP-CGG, at a 3:1 bead/cell ratio, and co-cultured for 4 days with OT-2 T cells (1:1 B/T cell ratio). FACS contour plots to the left show the appearance of a double positive (CD95^+^ GL7^+^) population in gated B220^+^ B cells. The bar plot to the right represents mean ± S.D. (*n* = 3). ** *p*<0.005; *** *p*<0.0005 (unpaired Student’s t test). **(B)** Proliferation of B cells from B1-8^hi^ transgenic WT and *Rhog*^−/−^ mice was calculated after 4 days of culture by CTV dilution. Data represent the mean ± S.D. (*n* = 3). * *p*<0.05 (unpaired Student’s t test). **(C)** Differentiation of naïve B cells from B1-8^hi^ transgenic WT and *Rhog*^−/−^ mice to GC B cells and plasma cells was followed along 7 days of in vitro culture with beads coated with either NIP-OVA or NP-CGG and OT-2 CD4^+^ T cells. The percentages of GC B cells were calculated according to CD38 downregulation and expression of intracellular Bcl-6 by gated B220^+^ B cells. The percentages of plasma cells were calculated according to the expression of CD138 and intracellular Blimp-1 by gated B220^+^ B cells. Line plots below represent mean ± S.D. (*n* = 3). **(D)** Expression of Bcl-6 and Blimp-1 by GC B cells was assessed by intracellular staining of B220^+^ B cells. Line plots below represent mean ± S.D. (*n* = 3). **(E)** Expression of Bcl-6 and Blimp-1 as a function of the number of cell divisions by naïve B cells from B1-8^hi^ transgenic WT mice stimulated 4 days in vitro with 1 μm beads coated with either NIP-OVA or NP-CGG and OT-2 T CD4^+^ T cells. The number of cell divisions was assessed by CTV dilution. Data represent mean ± S.D. (*n* = 3).

A key feature of GC B cell differentiation is the expression of the transcription factor Bcl-6 (Basso et al., 2012). We followed the emergence of a Bcl-6^+^ B cell population that also downregulates CD38 as GC markers for 7 days of WT B1-8^hi^ B cell culture with NIP-OVA-coated 1 μm beads and OT-2 T cells. We found a distinct Bcl-6^+^CD38^−^ population that reached a maximum of 25% of all B cells at day 5 of co-culture (Fig. 2C). GC B cell differentiation was also assessed using different combinations of GL7, CD95 and CD38, showing in this case that the maximum percentage of GC B cells was reached at day 4 to decay thereafter (Fig. 2C and Suppl. Fig. 1B). In parallel to GC markers we followed the expression of the plasmacytic B cell transcription factor Blimp-1 and the plasma cell marker CD138. Blimp-1 upregulation was detected at day 3 while the emergence of a distinct Blimp-1^+^CD138^+^ plasma cell population was clearly detected at day 4, reaching a plateau at day 5 (Fig. 2C). Bcl-6 and Blimp-1 are involved in a mutually regulatory loop in which Bcl-6 represses Blimp-1 and the latter represses Bcl-6, such that Blimp-1 expression favors the exit of cells from the GC differentiation program and terminal differentiation to plasma cells (Basso and Dalla-Favera, 2010; Rui et al., 2011). In fact, we found that Bcl-6 and Blimp-1 are expressed antagonistically in gated B220^+^GL7^+^ GC B cells in a way that by day 5 two distinct populations of GC B cells have clearly emerged: Bcl-6^high^Blimp-1^low^ and Bcl-6^low^Blimp-1^high^ (Fig. 2D). Analyzing Bcl-6 and Blimp-1 expression according to the number of B cell divisions at day 3 showed that Bcl-6 expression reached a maximum at the second cell division whereas Blimp-1 steadily increased up to the sixth cell division (Fig. 2E). The reduced Bcl-6 expression in highly divided cells could very well be originated by the growing repression exerted by Blimp-1 and responsible for the decline of Bcl-6 expression after 4 days of culture (Fig. 2D and 2E). These results indicate that B cell stimulation with a bead-bound antigen, which is taken up by phagocytosis, results in their differentiation to GC B cells in vitro that are regulated by Bcl-6 and Blimp-1 expression as it has been previously established in vivo (Basso et al., 2012; Rui et al., 2011).

To determine if OT-2 T cell proliferations and differentiation to TFH cells was also induced in co-culture with B1-8^hi^ B cells upon phagocytosis in an antigen-specific manner, we studied the expression of CXCR5, PD1 and ICOS markers of TFH cells in response to B1-8^hi^ B cells that had phagocytosed 1 or 3 μm beads coated with NIP covalently bound to OVA carrier protein (NIP-OVA, Suppl. Fig. 1C and 1D). Interestingly, proliferation of OT-2 T cells increased with the dose of beads whereas expression of surface markers was optimum at intermediate doses of beads that depended on their diameter. The bead-bound NIP-OVA stimulus also resulted in the generation of key TFH cell cytokines involved in the GC response —IL-4, IL-6, and IL-21—, that was inhibited if B1-8^hi^ B cells were deficient in RhoG (Suppl. Fig. 1E), strongly suggesting that B cell phagocytosis of antigen is required. Altogether, the above data suggest that B cells can phagocytose antigen and present it to cognate T cells that become activated and adopt markers and properties of TFH cells.

### Phagocytic B cells and T cells form organized structures in vitro

In germinal centers, antigen-specific B cells form clusters of highly proliferating cells that segregate from non-responding B cells in follicles. In the culture plates *in vit*ro, we found the formation of large clusters containing as many as 8,000 cells when B cells were stimulated with 1 μm beads coated with anti-IgM plus ovalbumin (Fig. 3A). The clusters consisted of a mixture of tightly intermingled B cells and CD4^+^ T cells. Similar clusters were found when mixtures of NP-specific B1-8^hi^ B cells and OT-2 T cells were incubated with 1 μm beads coated with NIP-OVA (Fig. 3B). Interestingly, stimulation of B cells with a similar dose (see below) of soluble NIP-OVA, resulted in the formation of much smaller clusters, suggesting that the large B cell and T cell aggregates were related to the phagocytic stimulus. Using mixtures of CTV-labeled B1-8^hi^ B cells and CFSE-labeled OT-2 T cells, after 7 days of stimulation with NIP-OVA antigen-coated beads we found that the periphery of the cluster contained the cells with most diluted CTV and CFSE, suggesting that B and T cells proliferate and expand towards the edges of the clusters (Fig. 3C). The periphery of the clusters also contained the highest percentage of B cells positive for the GC marker GL7 (Fig. 3D), suggesting that B cells proliferate and express GC markers towards the periphery. These data show that follicular B cells stimulated with antigen-coated beads form large clusters, together with T cells, that are reminiscent of germinal centers.

**Figure 3.**
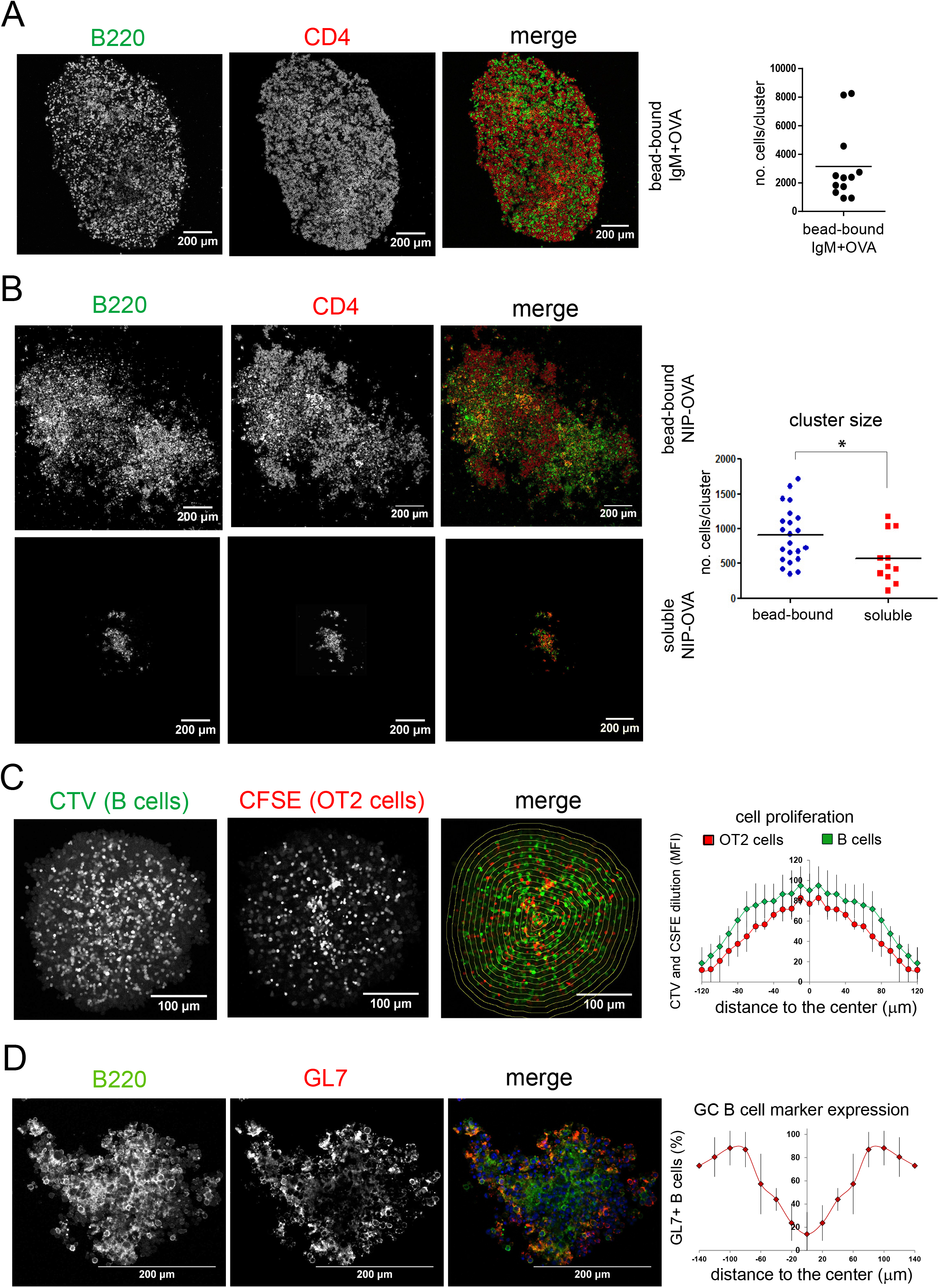
Phagocytic antigen induces the formation of large clusters of intermingled B and T cells. **(A)** Confocal microscopy image of a large cell cluster generated after 4 days of co-culture of OT-2 T cells and non-transgenic B cells stimulated with 1 μm beads coated with anti-IgM and OVA. B cells are stained with B220 in green; OT-2 T cells with CD4 in red. A quantification of the number of cells per cluster is shown in the plot to the right. **(B)** Confocal microscopy image of cell clusters generated after 7 days of incubation of naïve B cells from B1-8^hi^ transgenic WT mice with OT-2 T cells and either 1 μm beads coated with NIP-OVA or with soluble NIP-OVA. B220 is in green, CD4 in red. Cell number quantification per cluster for both antigen conditions is represented in the graph to the right. Data represent the mean ± S.D. * *p*<0.05 (unpaired Student’s t test). **(C)** Confocal microscopy image of a cluster of B1-8^hi^ transgenic WT B cells labeled with CTV and cultured for 7 days with CFSE-labeled OT-2 T cells and 1 μm beads coated with NIP-OVA. The intensity of CFSE and CTV staining was measured for all cells placed within the drawn concentric areas and represented in the plot to the right versus the distance to the center of the cluster. Data represent the mean ± S.D. for *n* = 5 clusters of similar size. **(D)** Confocal microscopy of a cluster generated by stimulation of B1-8^hi^ WT B cells with 1 μm beads coated with NIP-OVA for 7 days and stained with the B220 B cell marker (green) and the GL7 GC marker (red). GL7 intensity versus the distance to the center of the cluster was measured as in Fig. 3C. The line plot to the right represents the mean ± S.D. for *n* = 5 clusters of similar size.

### A bead-bound but not a soluble antigen induces the generation of class-switched immunoglobulins of high affinity

During the GC response, B cells recognize antigen through their BCR and this recognition has a dual effect: the activation of intracellular signaling pathways and antigen internalization for processing and presentation to CD4^+^ T cells. We wondered whether the way in which the antigen is presented to B cells (soluble versus phagocytic) influences B cell activation independently of antigen presentation to T cells. To normalize both stimuli for equal antigen presentation, we carried out a titration experiment in which proliferation of OVA-specific OT-2 CD4^+^ T cells was measured in response to B cells preincubated with different doses of soluble and bead-bound NIP-OVA. We found that a concentration of 100 ng/ml soluble NIP-OVA was as effective as a dose of 3 NIP-OVA-coated beads per B cell for inducing OT-2 cell proliferation (Fig. 4A). Furthermore, those doses of stimuli were equally effective at inducing the expression of Tfh cell markers in OT-2 T cells and T cell proliferation (Fig. 4B). In such conditions, B1-8^hi^ B cells differentiated into GC B cells independently of the soluble or bead-bound nature of the stimulus (Fig. 4C). However, soluble antigen was incompetent to promote the differentiation towards GC B cells that strongly bound NP antigen. These results suggest a difference between bead-bound and soluble stimuli not in the expression of GC B cell markers but on the selection of cells with higher affinity for antigen. In addition, the soluble antigen was significantly less mitogenic than the bead-bound one (Fig. 4D). As previously stated (Fig. 2A), RhoG deficiency impaired the formation of GC B cells in cultures of WT B1-8^hi^ B cells with bead-bound antigen (Fig. 4C). However, such deficiency was not apparent when antigen stimulus was given in soluble form, confirming that the bead-bound antigen needs to be phagocytosed by B cells in order to induce the formation of NP+ GC B cells. In addition to the different proliferation rate and NP-specific GC formation, bead-bound antigen was more effective at promoting expression of Bcl-6 and Blimp-1 than soluble antigen (Fig. 4E). By contrast, soluble antigen was more effective at inducing the expression of the cytidine deaminase (Aicda) mRNA than bead-bound antigen in spite the fact that bead-bound antigen was slightly more efficient at inducing somatic mutations in the nucleotide and protein sequences (Suppl. Fig. 2A). The mutation rates obtained in B1-8^hi^ B cells stimulated with bead-bound antigen were low but much higher than in OT-2 plus B cell cultures not stimulated with antigen. These results suggest that B cells with V region mutations are being selected for certain amino acid mutations in our *in vitro* system. Indeed, most of the missense mutations mapped at, or in the proximity of, the antigen-binding CDR sequences (Suppl. Fig. 2B). The effect of bead-bound antigen on upregulation of Aicda, Bcl-6 and Blimp1, and somatic mutation were related to phagocytic phenomena, since RhoG deficiency in B cells strongly reduced those processes (Fig. 4E and Suppl. Fig. 2A). These data indicate that a bead-bound, phagocytic-dependent antigen stimulus is more effective than soluble antigen at inducing B cell proliferation, as well as their differentiation into bona fide GC B cells. Indeed, bead-bound antigen was more effective than soluble antigen at inducing Ig class switch, which is a GC-dependent event, as evidenced by the expression of IgG1 at the B cell membrane (Fig. 4F). Furthermore, B cell stimulation with bead-bound antigen was a better inducer of the plasma cell differentiation marker CD138 than soluble antigen, suggesting that bead-bound but not soluble antigen induces B cell differentiation into antibody-secreting plasma cells (Fig. 4G).

**Figure 4.**
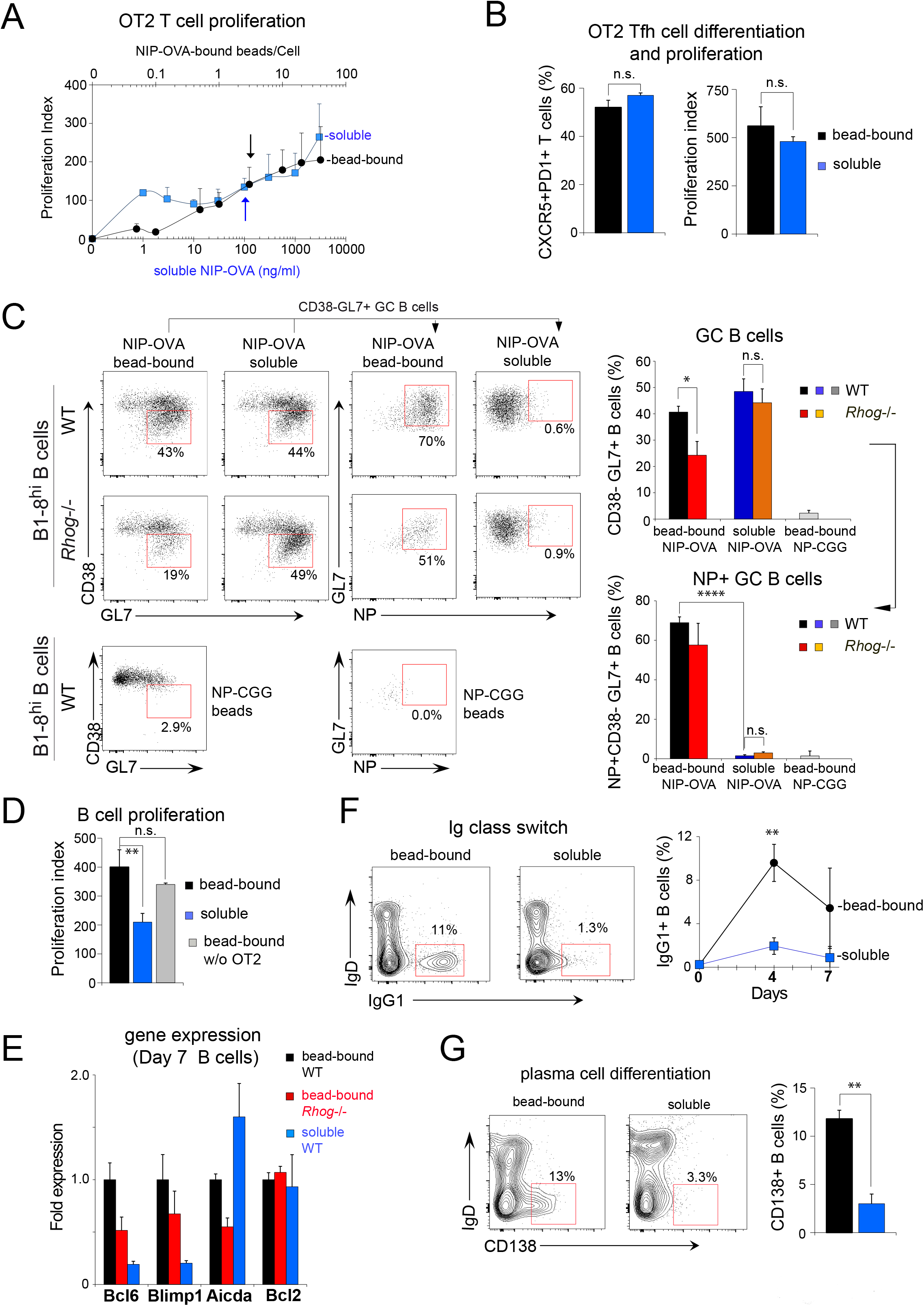
A phagocytic antigen is more efficient than a soluble one at inducing GC B cells in vitro. **(A)** Proliferation of OT-2 T cells after 4 days of culture with WT B1-8^hi^ B cells stimulated at different doses of bead-bound (black circles) or soluble (blue squares) NIP-OVA antigen. Arrows indicate the bead/B cell ratio (3: 1) and soluble antigen concentration (100 ng/ml) that induce comparable OT-2 T cell proliferation. **(B)** Graph plots of Tfh marker (CXCR5 and PD1) expression and proliferation of OT-2 T cells after 4 days of culture with the antigen conditions selected in (A). **(C)** Dot plots of GC marker expression by B cells from B1-8^hi^ transgenic WT and *Rhog*^−/−^ mice after 4 days of culture as in (A). A control of WT B1-8^hi^ B cells stimulated with a 3:1 ratio of NP-CGG-coated beads was cultured in parallel. The percentage of total GC B cells was calculated on B220^+^ B cells according to the expression of GL7 and CD38 (GL7^+^CD38^−^); the percentages of NP-specific GC B cells were calculated according to GL7 expression and NP binding (NP^hi^GL7^+^) on gated GL7^+^CD38^−^ B cells. Bar plots to the right show the percentages of total and NP+ GC B cells as mean ± S.D. (*n* = 3). * *p*<0.05; **** *p*<0.00005 (unpaired Student’s t test). **(D)** Proliferation of WT B1-8^hi^ B cells after 4 days of culture was calculated by CTV dilution after stimulation as in (A). A control of B cells cultured with bead-bound NIP-OVA in the absence of OT2 cells was carried out in parallel. Data represent the mean ± S.D. (*n* = 3). ** *p*<0.005 (unpaired Student’s t test). **(E)** RT-qPCR analysis of expression of the indicated genes performed on sorted WT and *Rhog*^−/−^ B1-8^hi^ B cells after 7 days of culture with OT-2 and specific stimulus as in (A). Bar plots show the fold induction mRNA expression of genes relative to the bead-bound WT condition. Expression of HPRT and 18S RNAs were used as normalizers. Data represent the mean ± S.D. (*n* = 3). **(F)** Contour plots showing the appearance of Ig class-switched IgG1^+^ IgD^−^ B1-8^hi^ B cells after 4 days in culture as in (A). The line plot to the right shows the appearance of IgG1^+^ B cells. Data represent the mean ± S.D. (*n* = 3). ** *p*<0.005 (unpaired Student’s t test). **(G)** Contour plots showing the appearance of plasma cells (B220^+^CD138^+^ IgD^−^) after 4 days in culture as in (A). Bar plots show the percentage of plasma cells (B220^int^ CD138^+^ IgD^−^) in those cultures. Data represent the mean ± S.D. (*n* = 3). ** *p*<0.005 (unpaired Student’s t test).

The capacity of bead-bound antigen to induce plasma cell differentiation was paralleled by the detection of high affinity anti-NP Igs in 7-day culture supernatants. The anti-NP Igs were comprised of non-switched IgM but also of high amounts of class-switched IgG1, IgG2a, IgG3, IgA and lesser, but detectable, amounts of IgG2b (Fig. 5A). The production of high and low affinity class-switched anti-NP Igs by bead-bound antigen-stimulated cells was strongly inhibited if B cells lacked RhoG, suggesting that their generation required the beads to be phagocytosed. Likewise, the supernatant of WT B1-8^hi^ B cell cultures stimulated with bead-bound antigen, but not with soluble antigen, contained high-affinity class-switched Igs, including IgG1, IgG2a, IgG2b, IgG3 and IgA (Fig. 5B). The above experiments were carried out with B1-8^hi^ B cells and OT-2 T cells purified by negative selection. To exclude the participation of a third cell type that could be contaminating the B and T cell populations, the experiments with bead-bound versus soluble NIP-OVA antigen were repeated with FACS-sorted follicular B (B220^+^CD23^+^CD43^−^CD11b^−^) and CD4^+^ OT-2 T cells (Suppl. Fig. 3A). In these conditions, B cells acquired GC markers (Suppl. Fig. 3B) and differentiated into antibody-producing cells able to secrete class-switched Igs (Suppl. Fig. 3C), thus indicating that T and B cells are sufficient to generate the GC-like reaction. Overall, these data showed that incubation of naïve B cells with a haptenated antigen immobilized onto 1 μm beads in the presence of antigen-specific helper CD4^+^ T cells *in vitro* results in the generation of high affinity antigen-specific antibodies of class-switched isotypes.

**Figure 5.**
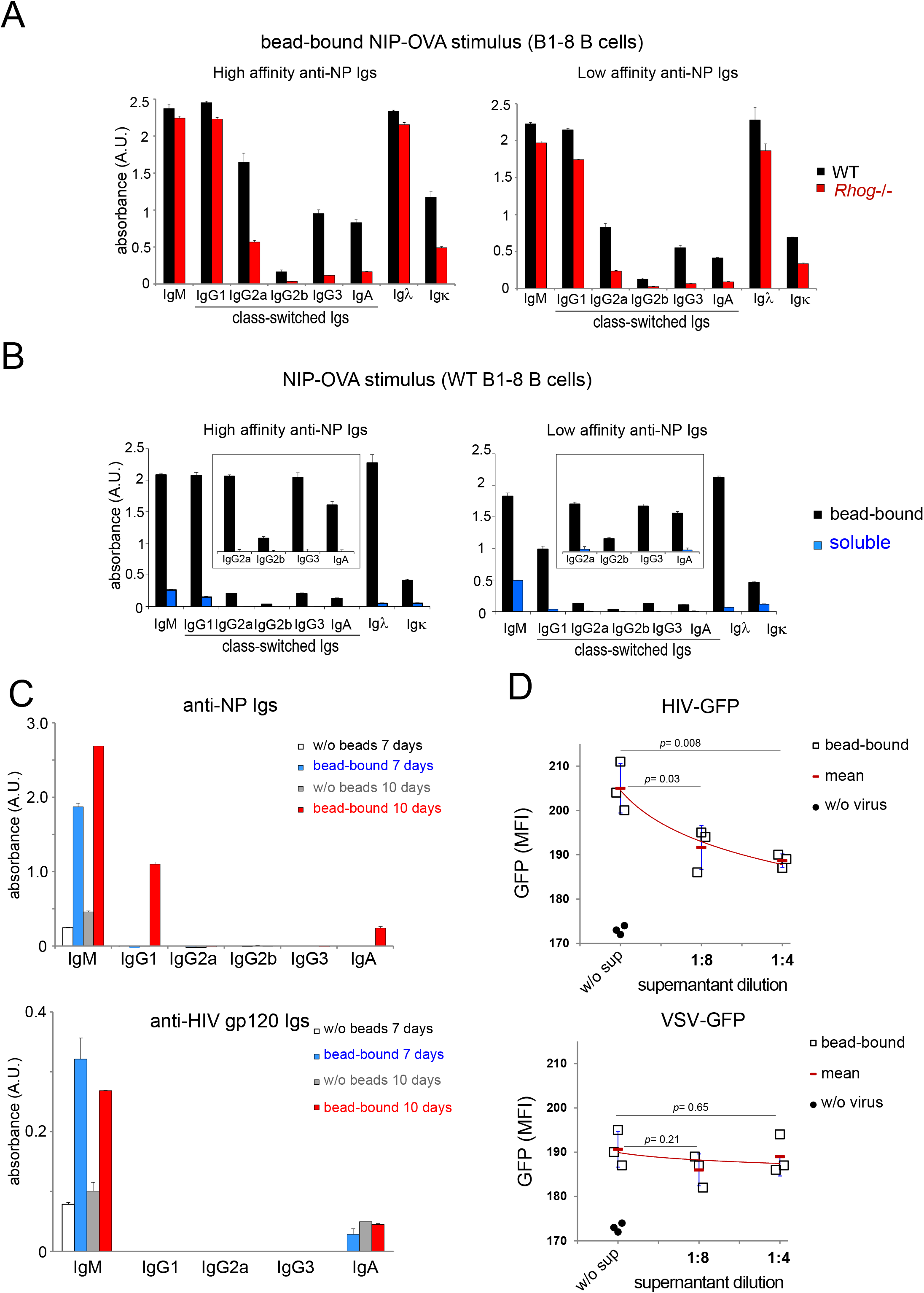
A phagocytic antigen induces the production of high-affinity class-switched and neutralizing antibodies in vitro. **(A)** Detection of high-affinity and low-affinity anti-NP Igs in supernatants from WT (black) or *Rhog*^−/−^ (red) B1-8^hi^ B cells cultured for 7 days with OT-2 T cells and bead-bound NIP-OVA (3:1 1 μm bead/B cell ratio). Data represent the mean ± S.D. (*n* = 3). **(B)** Detection of high- and low-afinity anti-NP Igs in supernatants of B1-8^hi^ B cells stimulated with soluble (blue) or bead-bound (black) NIP-OVA together with OT-2 T cells for 7 days. Data represent the mean ± S.D. (*n* = 3). **(C)** Detection of high affinity anti-NP Igs and anti-HIV Env protein Igs in culture supernatants of non-transgenic B cells stimulated with 1 μm beads (3:1 ratio) coated with NIP-OVA and HIV Env recombinant protein together with OT-2 T cells for 7 days or 10 days. 10-day cell cultures were supplemented at day 5 with 1 ng/ml IL-4 and 10 ng/ml IL-21. Data represent the mean ± S.D. (*n* = 3). **(D)** Presence of HIV neutralizing antibodies in the culture supernatants of (C) manifested as the inhibition of entry of GFP-expressing HIV virions coated with either the HIV Env protein or pseudotyped with the VSV G protein in MOLT-4 T cells. Data represent the mean ± S.D. (*n* = 3). *p* values were assessed using an unpaired Student’s t test.

### B cell stimulation with bead-bound antigen generates functional antibodies from non-transgenic B cells *in vitro*

To interrogate if the phagocytic-dependent antigen delivery to B cells can be used to generate antigen-specific antibodies in vitro out of a non-transgenic BCR repertoire, we incubated NP-OVA-coated 1 μm beads with purified B cells from non-transgenic C57BL/6 mice and with OT-2 T cells. After 7 days, the culture supernatant contained anti-NP IgMs but not detectable class-switched Igs (Fig. 5C). Since the concentrations of IL-4 and IL-21 in the culture supernatants of bead-bound antigen-stimulated B cells was low (Suppl. Fig. 1E) we attempted to extend the life of the cultures by adding recombinant IL-4 and IL-21 at day 5. We found detectable anti-NP IgG1 and IgA in the cytokine-supplemented cultures after 10 days of incubation (Fig. 5C).

To determine if the *in vitro* system can be used to generate antibodies of medical interest, we coated 1 μm beads with Env recombinant protein of HIV-1 and NIP-OVA as carrier protein. Coated beads were incubated with B cells from non-transgenic C57BL/6 mice and OT-2 T cells. We detected the generation of Env-specific IgMs both at day 7 and day 10 but not class-switched Igs (Fig. 5C). Nonetheless, the supernatant of 7-day cultures was tested for its capacity to inhibit the entry of HIV-GFP viral particles coated with the HIV-1 Env protein or pseudotyped with the envelope protein of VSV. The supernatant dose-dependently inhibited the entry of the HIV Env-mediated virus but not of the VSV GFP-dependent one, suggesting that the generated anti-HIV Env IgMs specifically neutralize HIV.

### A bead-bound phagocytic stimulus provides a strong and sustained BCR signal

To provide a mechanistic explanation to the above findings we interrogated if the bead-bound stimulus could result in a more intense or more sustained BCR signal than soluble antigen. We first determined the degree of occupancy of the BCR in both conditions using a fluorescent NP derivative. At the conditions used above for comparison (3 coated beads vs. 100 ng/ml of soluble protein), 35% of the B1-8 BCR was free to bind NP hapten in cells incubated with beads, whereas only 1% was free if cells had been incubated with soluble protein (Fig. 6A). We also measured BCR downregulation as a function of time and found that the soluble stimulus was at least as effective as the bead-bound stimulus at promoting BCR downregulation (Fig. 6B). These data indicate that the bead-bound antigen is not more effective than the soluble one in terms of BCR occupancy or BCR downregulation. Therefore, its superior capacity to produce class-switched high affinity antigen-specific Igs is not explained by simply higher BCR occupation. We therefore investigated if signaling events downstream of the BCR were differentially activated. Phagocytosis requires the rearrangement of the actin cytoskeleton around the particle in the phagocytic cup (Yuseff et al., 2013). We thus investigated if there were differences in terms of the intensity or duration of actin polymerization in B cells incubated with bead-bound vs. soluble antigen. Both stimuli equally increased polymerized F-actin levels in B cells after 1 min of incubation (Fig. 6C). However, whereas the polymerization phase was rapidly followed by an intense depolymerization in B cells stimulated with soluble antigen, the high F-actin content was sustained in B cells stimulated with bead-bound antigen. The bead-bound stimulus elicited a more intense and sustained phosphorylation of Akt and ERK, which are two events linked to activation of the PI3K and Ras pathways, than the soluble stimulus (Fig. 6D). More importantly, phosphorylation of Syk, a direct BCR effector previously shown to mediate FcγR- and CR-dependent phagocytosis (Shi et al., 2006; Tohyama and Yamamura, 2006), was also more intense and sustained (Fig. 6D). This suggests that the bead-bound stimulus induces a stronger BCR signal that is more persistent in time than the soluble stimulus. To determine if the stronger signal promoted by the bead-bound stimulus was related to the phagocytic process, we compared the phosphorylation of Akt and S6 (in the PI3K pathway) and of ERK in WT vs RhoG-deficient B cells in response to bead-bound antigen. We found that RhoG is required to induce and sustain those signals, as well as the phosphorylation of Syk and of the Igα subunit of the BCR, strongly suggesting that antigen phagocytosis elicits a longer and more intense BCR signal (Fig. 6E).

**Figure 6.**
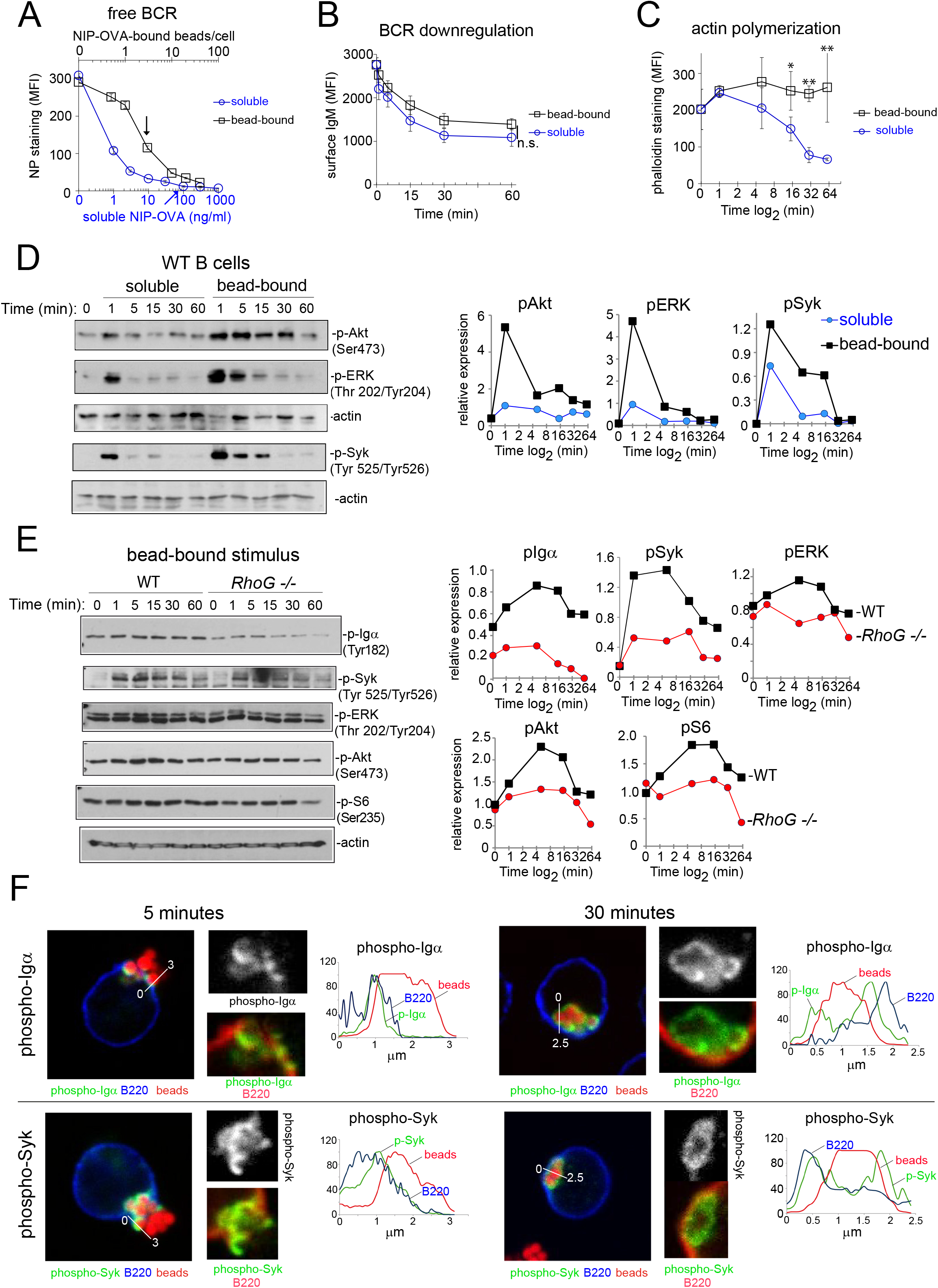
A phagocytic antigen induces a stronger and more sustained BCR signal than a soluble one. **(A)** Surface BCR saturation plot of purified B1-8^hi^ B cells incubated with bead-bound (black) or soluble NIP-OVA (blue) antigen at different doses for 1 hour at 0°C. Free unbound BCR was estimated by staining with NP-PE. Data represent the mean ± S.D. (*n* = 3). Arrows indicate the bead-dose and soluble concentration determined previously with comparable effects on OT-2 T cell proliferation (Fig. 4A). **(B)** BCR downmodulation was estimated according to anti-IgM staining of B1-8^hi^ B cells after stimulation with bead-bound (red, 3:1 ratio) or soluble (blue, 100 ng/ml) NIP-OVA antigen for different time-points at 37°C. Data represent the mean ± S.D. (*n* = 3). **(C)** F-actin content was measured by phalloidin staining of B1-8^hi^ B cells after stimulation with bead-bound (black, 3:1 ratio) or soluble (blue, 100 ng/ml) NIP-OVA antigen for different time-points at 37°C. Data represent the mean ± S.D. (*n* = 3). * *p*<0.05; ** *p*<0.005 (unpaired Student’s t test). **(D)** Immunoblot analysis of phosphorylation events downstream of the BCR after stimulation of WT B1-8^hi^ B cells with either a 3:1 ratio of bead-bound NIP-OVA or with 100ng/ml of soluble NIP-OVA for different time-points. Plots to the right show protein phosphorylation levels relative to the amount of actin quantified by densitometry. **(E)** Immunoblot analysis of phosphorylation events downstream of the BCR after stimulation of WT (red) or RhoG-deficient (blue) B1-8^hi^ B cells with a 3:1 ratio of bead-bound NIP-OVA for different time points. Plots to the right show protein phosphorylation levels relative to the amount of actin quantified by densitometry. **(F)** Midplane confocal microscopy images of B1-8^hi^ B cells in the process of phagocytosing (5 min. of incubation) or having completely phagocytosed (30 min.) 1 μm beads coated with NIP-OVA. Details of the phagocytic cups (5 min) and the phagosomes (30 min.) are shown in the enlarged pictures. The B220 B cell marker is in blue, beads in red and phospho-Igα and phospho-Syk antibodies in green. Histogram overlays show the signal intensity in the 3 colors along the white lines drawn in the main images.

We next assessed the cellular location of phosphorylated BCRs during antigen phagocytosis. Using fluorescent 1 μm beads and confocal microscopy we found that in B cells stimulated with bead-bound antigen for a short time (5 min), both phospho-Igα and phospho-Syk were only found in the phagocytic cups (Fig. 6F). Interestingly, both proteins were found to still be phosphorylated all around the phagocytosed beads at a late (30 min) time point. These results show that BCR phosphorylation persists in the intracellular phagosome and suggest that this might be the cause for sustained BCR signaling when antigen is taken up by a phagocytic mechanism.

## DISCUSSION

In this study we describe a system to recreate a GC in vitro based on the use of only two cell types, B cells and CD4+ T cells, and on the administration of antigen to B cells associated to a particle that needs to be phagocytosed by a BCR-mediated mechanism. Unlike previously described methods (Nojima et al., 2011), the phagocytosis-based method allows to recreate a GC in an antigen-specific manner and can therefore be used both to study T-B cell interactions during GC formation and to select B cells that produce high-affinity class switched antibodies of therapeutic or diagnostic interest. We show that B cells that have phagocytosed large inert particles 1 μm coated either with an antibody that triggers the BCR or with its specific antigen, can efficiently present antigen derived from these particles to CD4^+^ T cells *in vitro.* In exchange, T cells proliferate and express markers of differentiation towards TFH cells. In turn, TFH cells provide help to the antigen-presenting B cells and favor their differentiation into GC B cells, promoting Ig class switch and somatic mutation, two key features of a mature GC-derived humoral response. Indeed, using this simple two cell-type system, we detect the production of antigen-specific high affinity antibodies of all isotypes, resulting in a method to recreate antigen-specific GCs *in vitro*. Key to this success is the acquisition of antigen through a BCR-dependent phagocytic process linked to stronger and more sustained BCR signaling compared to a soluble antigen stimulus. During phagocytosis, BCR signaling initiates in the phagocytic cup and persists once the particle has been completely phagocytosed. The phagocytic process is mediated in part by the GTPase RhoG, which has been previously involved in the phagocytosis of apoptotic bodies by macrophages, and in both type I and type II phagocytosis (Niedergang et al., 2016; Tzircotis et al., 2011). Although the defect in RhoG does not lead to a blockade of B cell phagocytosis and GC formation *in vitro*, perhaps because of a possible redundancy with other Rho GTPases (Tzircotis et al., 2011), we previously showed that RhoG-deficient B cells are a valuable tool to determine the role of antigen phagocytosis by B cells in the humoral response *in vivo* (Martinez-Riano et al., 2018). Although it has been previously shown that B1 cells and to a lesser extent follicular B2 cells can phagocytose bacteria (Gao et al., 2012; Plzakova et al., 2015; Zhu et al., 2016), the prevalent view is that B cells acquire antigen from immune complexes retained at the surface of FDCs or macrophages (Avalos and Ploegh, 2014; Suzuki et al., 2009). Although our data do not oppose this view, they highlight the relevance of the phagocytic process for GC differentiation. Indeed, it has been shown that antigen is captured by binding to the BCR in a process that also physically extracts membrane components from the cell that presents antigen to the B cell (Batista et al., 2001). This phenomenon is very much reminiscent of the trogocytosis process by which T cells acquire MHC and membrane fragments from APC and that we previously characterized as a phagocytic process mechanistically mediated by RhoG (Martinez-Martin et al., 2011). Therefore, it could very well be that the mechanism of acquisition of antigen by B cells from FDCs and macrophages displaying immune complexes is indeed phagocytic. Still, what is the relevance of antigen phagocytosis by B cells in the GC response? It has been shown that the endocytosed BCR continues signaling from intracellular compartments, which is important to sustain the activation of the PI3K-Akt pathway (Chaturvedi et al., 2011). We show here that phagocytosis promoted by the BCR results in a stronger and more sustained phosphorylation of the BCR and downstream targets (including Akt) than a soluble stimulus, presumably internalized by B cells via an endocytic mechanism. Phagocytosis can also be a mechanism for selection of B cells bearing BCRs with higher affinity for antigen. High affinity for antigen correlates with enhanced CD4^+^ T cell activation (Batista and Neuberger, 1998) and mechanical forces are used to discriminate between antigen affinities by B cells (Natkanski et al., 2013). Interestingly, antigen uptake by B cells from artificial membranes used in the latter study require myosin II and this myosin is known to be required for phagocytic cup squeezing (Araki, 2006). Hence, the energetic requirements for phagocytosis may make this mechanism of antigen uptake by B cells a discriminatory sensor of BCR affinity.

Regardless of the involvement of a phagocytic process in antigen uptake by B cells from antigen-presenting FDCs or macrophages, the direct phagocytosis of pathogens or particles mediated by the BCR may be relevant during the physiological humoral response to infections and/or vaccination. Indeed, it has been reported that mice with impaired antigen presentation by FDC still generate mature Igs when immunized with adjuvants, suggesting that antigen acquisition by germinal center B cells must involve additional processes (Chen et al., 2000; Wu et al., 2000). It has long been known that B cells can acquire antigen and present it to CD4^+^ T cells by a phagocytic process (Vidard et al., 1996). However, to our knowledge its relevance and possible consequence for GC formation has not been previously described. We show that an improvement in the humoral response promoted by the combination of a haptenated antigen with alum adjuvant is completely dependent on the capacity of B cells to phagocytose antigen. Although known for a long time to benefit the humoral response and used as a component of vaccines involving recombinant proteins, the mechanism by which alum provides this help is still under debate. Since alum makes aggregates of 1-10 μm with antigen (Lindblad, 2004), which are in the range of sizes that can be phagocytosed by B cells, we propose that at least one of the mechanisms by which alum is an effective adjuvant is by favoring the acquisition of antigen by B cells by a phagocytic mechanism. To what extent direct phagocytosis of bacterial or fungal pathogens by B cells is involved in the mature humoral response against those pathogens is a pending issue.

In summary, our study shows a method to recreate antigen-specific GC responses in vitro that can be used both to mechanistically study T cell: B cell interaction requirements for GC responses in detail and to select and generate high-affinity class-switched immunoglobulins of clinical or diagnostic interest in vitro. In addition, our study highlights the relevance that antigen phagocytosis by B cells may have for the humoral response to pathogens and vaccines and provides a mechanistic explanation based on the intensity and duration of the BCR signal.

## EXPERIMENTAL PROCEDURES

### Mice

*Rras2*^−/−^ and *Rhog*^−/−^ mice were generated as previously described (Delgado et al., 2009; Vigorito et al., 2004). These mice were crossed with NP-specific B1-8^hi^ knockin mice bearing a pre-rearranged V region (Shih et al., 2002). Mice transgenic for the OT-2 TCR specific for a peptide 323-339 of chicken ovoalbumin presented by I-A^b^ (Barnden et al., 1998) and C57BL/6 bearing the pan-leukocyte marker allele CD45.1 were kindly provided by Dr. Carlos Ardavín (CNB, Madrid). All animals were backcrossed to the C57BL/6 background for at least 10 generations. For all in vivo experiments, age (6-10 weeks) and sex were matched between the *Rhog*^+/+^ (WT) and *Rhog*^−/−^ mice. Mice were maintained under SPF conditions in the animal facility of the Centro de Biología Molecular Severo Ochoa in accordance with applicable national and European guidelines. All animal procedures were approved by the ethical committee of the Centro de Biología Molecular Severo Ochoa.

### Antigen-coated bead preparation

To prepare beads with adsorbed antigen, a total of 130×10^6^carboxylated latex beads of 1 μm diameter were incubated overnight with a concentration of 40 μg/ml of protein in 1 ml of PBS at 4°C. For preparation of antigen-coated beads of 3 and 10 μm diameter, bead concentration was reduced in a staggered way; 3-fold and 30-fold, respectively. Beads were subsequently washed twice with PBS plus 1% BSA and resuspended in RPMI medium. To prepare beads with covalently-bound antigen, the PolyLink Protein Coupling Kit (Polysciences) was used as indicated by the manufacturer. An equivalent of 12.5 mg of beads were washed in Coupling Buffer (50 mM MES, pH 5.2, 0.05% Proclin 300), centrifuged 10 minutes at 1000g and resuspended in 170 μL Coupling Buffer. A 20 μl volume of Carbodiimide solution (freshly prepared at 200 mg/ml) was added to the bead suspension and incubated for 15 minutes. After that, a total of 400 μg of NIP-OVA were added at a concentration of 5 mg/ml final concentration.

To prepare beads coupled to two different proteins we followed a sequential procedure: the first protein was added at sub-saturating conditions (100 μg p17/p24/gp120 HIV-1 protein) for one hour and after that the second one was added to reach saturation (400 μg NIP-OVA) and incubated one additional hour. Incubations were carried out at room temperature with gentle mixing. Beads were centrifuged and washed twice in Wash/Storage buffer (10 mM Tris, pH 8.0, 0.05% BSA, 0.05% Proclin 300). To remove non-covalent bound protein, beads were washed once with 0.1% SDS followed by two washes with PBS + 1% BSA for SDS removal.

### Proliferation and stimulation assays

Proliferation of OT-II and B cells was assessed using CFSE or CellTrace Violet (CTV) labelling as specified by the manufacturer (Thermofisher). A total of 2×10^5^ purified naïve B cells were CTV-stained and co-cultured with purified CFSE-stained OT-II T cells at a 1:1 ratio together with antigen-coated beads or soluble antigen in a round-bottom 96-well plate. For the bead-bound stimulus, B cells were incubated with 1 μm beads coated with NIP-OVA, NP-CGG or anti-IgM plus ovalbumin at different bead:B cell ratios. For stimulation with soluble antigen, NIP-OVA was used at a concentration of 100 ng/ml. After 3-4 days of culture, cells were washed in PBS plus 1% BSA and stained for T cell activation (CD25, CD44), TFH (CXCR5, PD1, ICOS) or germinal center B cell (CD95, GL7, CD38) markers. To study differentiation of these cultured B cells to plasma cells, the cells were left in culture for 4 and 7 days and stained for CD138, IgD, and IgG1. The intracellular stainings for Bcl-6 and Blimp1 were performed using the Foxp3/Transcription Factor Staining Buffer Set. Samples were analysed by FACS (FACS Canto II) and FlowJo software.

### Measurement of antigen-specific antibodies

To measure the release of NP-specific Ig in vitro, B cell: OT-2 T cell culture supernatants were incubated on NP(7)-BSA-coated or NP(41)-BSA coated Costar p96 flat-bottom plates to measure the release of high- and low-affinity Igs, respectively. The SBA Clonotyping System-HRP (Southern Biotech) was used to detect the presence of antigen-specific Ig isotypes. When B1-8^hi^ transgenic B cells were used, purified B2 B cells and OT-2 T cells were cultured at 1:1 ratio for 7 days in the presence of NIP-OVA-coated 1 μm beads (3:1 bead/B cell ratio) or 100ng/ml soluble NIP-OVA. For cultures of non-transgenic B cells, purified naïve C57BL/6 B2 cells were preincubated with a mixture of NIP-OVA and HIV-1 p17/p24/gp120 fusion protein (Jena Biosciences) covalently bound to 1 μm beads (1:1 bead:Bcell ratio) and cultured with OT-2 T cells (1:1 B cell/OT-2 T cell ratio). After 5 days of culture, some cultures were supplemented with 1 ng/mL IL-4 and 10 ng/mL IL-21 (Peprotech). Supernatants were recovered at day 10 and used to measure Igs by ELISA.

### HIV neutralization assay

Lentiviral supernatants were produced from transfected HEK-293T cells as described previously (Martinez-Martin et al., 2009). Briefly, lentivirus were obtained by co-transfecting plasmids psPAX2 (gag/pol), pHRSIN-GFP and either HIV-1 envelope (pCMV-NL4.3) or VSV envelope (pMD2.G) using the JetPEI transfection reagent (Polyplus Transfection). Viral supernatants were obtained after 24 and 48 hours of transfection. Polybrene (8μg/mL) was added to the viral supernatants prior to transduction of MOLT-4 cells. A total of 3×10^5^ MOLT-4 cells were plated on a P24 flat-bottom well 350 μL of DMEM and 350 μL of viral supernatant were added. Cells were centrifuged for 90 minutes at 2200 rpm and left in culture for 24 hours.

The culture supernatants of purified naïve C57BL/6 B2 cells stimulated with a mixture of NIP-OVA and HIV-1 p17/p24/gp120 fusion protein covalently bound to 1 μm beads (1 bead:Bcell ratio) together with OT-2 T cells (1:1 B cell/OT-2 T cell ratio) were incubated at different dilutions (1:8 and 1:4) with the viral supernatant for 1 hour at 37 °C. This mixture was subsequently used to infect MOLT-4 cells. As a control of infectivity, MOLT-4 cells were infected with viral supernatant without antibody supernatants. MOLT-4 cell infection was assessed according to GFP expression by Flow Cytometry (FACS Canto II).

## QUANTIFICATION AND STATISTICAL ANALYSIS

### Statistical analysis

Statistical parameters including the exact value of n, the means +/− s.d. are reported in the Figure and Figure legends. A non-parametric two-tailed unpaired t-test was used to assess the confidence intervals.

## Acknowledgments

We thank Nuria Martínez-Martín and Facundo Batista for critical reading of the manuscript. We are also indebted to Cristina Prieto, Valentina Blanco, and Tania Gómez for their expert technical assistance and Belen de Andrés and Maria Angeles Muñoz-Alcalá for helpful comments and advice on methodology.

## Author contributions

Conceptualization, A. M.-R. P.D. and B.A.; Methodology, A. M.-R., E. R.-B. and P.M.; Software, D.A.; Formal Analysis, B.A.; Investigation, A. M.-R., P.M. and E.R.-B.; Writing-Original Draft, B.A.; Writing-Review & Editing, A. M.-R., C.L.O. and M.T.; Funding Acquisition, B.A. and M.T.; Resources, B.A. and M.T.; Supervision, B.A.

## REFERENCES

Araki, N. (2006). Role of microtubules and myosins in Fc gamma receptor-mediated phagocytosis. Front Biosci. 11, 1479–1490.

Avalos, A.M., and Ploegh, H.L. (2014). Early BCR Events and Antigen Capture, Processing, and Loading on MHC Class II on B Cells. Front Immunol. 5:92., 10.3389/fimmu.2014.00092. eCollection 02014.

Barnden, M.J., Allison, J., Heath, W.R., and Carbone, F.R. (1998). Defective TCR expression in transgenic mice constructed using cDNA-based alpha- and beta-chain genes under the control of heterologous regulatory elements. Immunol Cell Biol. 76, 34–40.

Basso, K., and Dalla-Favera, R. (2010). BCL6: master regulator of the germinal center reaction and key oncogene in B cell lymphomagenesis. Adv Immunol. 105:193–210., 10.1016/S0065-2776(1010)05007-05008.

Basso, K., Dalla-Favera, R., Cattoretti, G., Angelin-Duclos, C., Shaknovich, R., Zhou, H., Wang, D., and Alobeid, B. (2012). Roles of BCL6 in normal and transformed germinal center B cells PRDM1/Blimp-1 is expressed in human B-lymphocytes committed to the plasma cell lineage. Immunol Rev. 247, 172–183. doi: 110.1111/j.1600-1065X.2012.01112.x.

Batista, F.D., Iber, D., and Neuberger, M.S. (2001). B cells acquire antigen from target cells after synapse formation. Nature. 411, 489–494.

Batista, F.D., and Neuberger, M.S. (1998). Affinity dependence of the B cell response to antigen: a threshold, a ceiling, and the importance of off-rate. Immunity. 8, 751–759.

Chaturvedi, A., Martz, R., Dorward, D., Waisberg, M., and Pierce, S.K. (2011). Endocytosed BCRs sequentially regulate MAPK and Akt signaling pathways from intracellular compartments. Nat Immunol. 12, 1119–1126. doi: 1110.1038/ni.2116.

Chen, Z., Koralov, S.B., Gendelman, M., Carroll, M.C., and Kelsoe, G. (2000). Humoral immune responses in Cr2-/-mice: enhanced affinity maturation but impaired antibody persistence. Journal of immunology 164, 4522–4532.

deBakker, C.D., Haney, L.B., Kinchen, J.M., Grimsley, C., Lu, M., Klingele, D., Hsu, P.K., Chou, B.K., Cheng, L.C., Blangy, A., et al. (2004). Phagocytosis of apoptotic cells is regulated by a UNC-73/TRIO-MIG-2/RhoG signaling module and armadillo repeats of CED-12/ELMO. Curr Biol. 14, 2208–2216.

Delgado, P., Cubelos, B., Calleja, E., Martinez-Martin, N., Cipres, A., Merida, I., Bellas, C., Bustelo, X.R., and Alarcon, B. (2009). Essential function for the GTPase TC21 in homeostatic antigen receptor signaling. Nat Immunol. 10, 880–888. doi: 810.1038/ni.1749. Epub 2009 Jun 1028.

Gao, J., Ma, X., Gu, W., Fu, M., An, J., Xing, Y., Gao, T., Li, W., and Liu, Y. (2012). Novel functions of murine B1 cells: active phagocytic and microbicidal abilities. Eur J Immunol. 42, 982–992. doi: 910.1002/eji.201141519.

Gitlin, A.D., Shulman, Z., and Nussenzweig, M.C. (2014). Clonal selection in the germinal centre by regulated proliferation and hypermutation. Nature. 509, 637–640. doi: 610.1038/nature13300. Epub 12014 May 13304.

Groves, E., Dart, A.E., Covarelli, V., and Caron, E. (2008). Molecular mechanisms of phagocytic uptake in mammalian cells. Cell Mol Life Sci. 65, 1957–1976.

Henson, P.M. (2005). Engulfment: ingestion and migration with Rac, Rho and TRIO. Curr Biol. 15, R29–30.

Li, J., Barreda, D.R., Zhang, Y.A., Boshra, H., Gelman, A.E., Lapatra, S., Tort, L., and Sunyer, J.O. (2006). B lymphocytes from early vertebrates have potent phagocytic and microbicidal abilities. Nat Immunol. 7, 1116–1124. Epub 2006 Sep 1117.

Lindblad, E.B. (2004). Aluminium compounds for use in vaccines. Immunol Cell Biol. 82, 497–505.

Martinez-Martin, N., Fernandez-Arenas, E., Cemerski, S., Delgado, P., Turner, M., Heuser, J., Irvine, D.J., Huang, B., Bustelo, X.R., Shaw, A., and Alarcon, B. (2011). T cell receptor internalization from the immunological synapse is mediated by TC21 and RhoG GTPase-dependent phagocytosis. Immunity. 35, 208–222. Epub 2011 Aug 2014.

Martinez-Martin, N., Risueno, R.M., Morreale, A., Zaldivar, I., Fernandez-Arenas, E., Herranz, F., Ortiz, A.R., and Alarcon, B. (2009). Cooperativity between T cell receptor complexes revealed by conformational mutants of CD3epsilon. Sci Signal. 2, ra43.

Martinez-Riano, A., Bovolenta, E.R., Mendoza, P., Oeste, C.L., Martin-Bermejo, M.J., Bovolenta, P., Turner, M., Martinez-Martin, N., and Alarcon, B. (2018). Antigen phagocytosis by B cells is required for a potent humoral response. EMBO Rep 19.

Nakashima, M., Kinoshita, M., Nakashima, H., Habu, Y., Miyazaki, H., Shono, S., Hiroi, S., Shinomiya, N., Nakanishi, K., and Seki, S. (2012). Pivotal advance: characterization of mouse liver phagocytic B cells in innate immunity. J Leukoc Biol. 91, 537–546. doi: 510.1189/jlb.0411214. Epub 0412011 Nov 0411214.

Natkanski, E., Lee, W.Y., Mistry, B., Casal, A., Molloy, J.E., and Tolar, P. (2013). B cells use mechanical energy to discriminate antigen affinities. Science. 340, 1587–1590. doi: 1510.1126/science.1237572. Epub 1232013 May 1237516.

Niedergang, F., Di Bartolo, V., and Alcover, A. (2016). Comparative Anatomy of Phagocytic and Immunological Synapses. Front Immunol. 7:18., 10.3389/fimmu.2016.00018. eCollection 02016.

Nojima, T., Haniuda, K., Moutai, T., Matsudaira, M., Mizokawa, S., Shiratori, I., Azuma, T., and Kitamura, D. (2011). In-vitro derived germinal centre B cells differentially generate memory B or plasma cells in vivo. Nat Commun. 2:465., 10.1038/ncomms1475.

Nowosad, C.R., Spillane, K.M., and Tolar, P. (2016). Germinal center B cells recognize antigen through a specialized immune synapse architecture. Nat Immunol. 17, 870–877. doi: 810.1038/ni.3458. Epub 2016 May 1016.

Ochando, J.C., Homma, C., Yang, Y., Hidalgo, A., Garin, A., Tacke, F., Angeli, V., Li, Y., Boros, P., Ding, Y., et al. (2006). Alloantigen-presenting plasmacytoid dendritic cells mediate tolerance to vascularized grafts. Nat Immunol. 7, 652–662. Epub 2006 Apr 2023.

Parra, D., Rieger, A.M., Li, J., Zhang, Y.A., Randall, L.M., Hunter, C.A., Barreda, D.R., and Sunyer, J.O. (2012). Pivotal advance: peritoneal cavity B-1 B cells have phagocytic and microbicidal capacities and present phagocytosed antigen to CD4+ T cells. J Leukoc Biol. 91, 525–536. doi: 510.1189/jlb.0711372. Epub 0712011 Nov 0711374.

Phan, T.G., Green, J.A., Gray, E.E., Xu, Y., and Cyster, J.G. (2009). Immune complex relay by subcapsular sinus macrophages and noncognate B cells drives antibody affinity maturation. Nat Immunol. 10, 786–793. doi: 710.1038/ni.1745. Epub 2009 Jun 1037.

Plzakova, L., Krocova, Z., Kubelkova, K., and Macela, A. (2015). Entry of Francisella tularensis into Murine B Cells: The Role of B Cell Receptors and Complement Receptors. PLoS ONE. 10, e0132571. doi: 0132510.0131371/journal.pone.0132571. eCollection 0132015.

Ramiscal, R.R., and Vinuesa, C.G. (2013). T-cell subsets in the germinal center. Immunol Rev. 252, 146–155. doi: 110.1111/imr.12031.

Rui, L., Schmitz, R., Ceribelli, M., and Staudt, L.M. (2011). Malignant pirates of the immune system. Nat Immunol. 12, 933–940. doi: 910.1038/ni.2094.

Shi, Y., Tohyama, Y., Kadono, T., He, J., Miah, S.M., Hazama, R., Tanaka, C., Tohyama, K., and Yamamura, H. (2006). Protein-tyrosine kinase Syk is required for pathogen engulfment in complement-mediated phagocytosis. Blood 107, 4554–4562.

Shih, T.A., Roederer, M., and Nussenzweig, M.C. (2002). Role of antigen receptor affinity in T cell-independent antibody responses in vivo. Nat Immunol. 3, 399–406. Epub 2002 Mar 2018.

Suzuki, K., Grigorova, I., Phan, T.G., Kelly, L.M., and Cyster, J.G. (2009). Visualizing B cell capture of cognate antigen from follicular dendritic cells. J Exp Med. 206, 1485–1493. doi: 1410.1084/jem.20090209. Epub 20092009 Jun 20090208.

Tohyama, Y., and Yamamura, H. (2006). Complement-mediated phagocytosis--the role of Syk. IUBMB life 58, 304–308.

Tzircotis, G., Braga, V.M., and Caron, E. (2011). RhoG is required for both FcgammaR- and CR3-mediated phagocytosis. J Cell Sci. 124, 2897–2902.

Vidard, L., Kovacsovics-Bankowski, M., Kraeft, S.K., Chen, L.B., Benacerraf, B., and Rock, K.L. (1996). Analysis of MHC class II presentation of particulate antigens of B lymphocytes. J Immunol. 156, 2809–2818.

Vigorito, E., Bell, S., Hebeis, B.J., Reynolds, H., McAdam, S., Emson, P.C., McKenzie, A., and Turner, M. (2004). Immunological function in mice lacking the Rac-related GTPase RhoG. Mol Cell Biol. 24, 719–729.

Wu, X., Jiang, N., Fang, Y.F., Xu, C., Mao, D., Singh, J., Fu, Y.X., and Molina, H. (2000). Impaired affinity maturation in Cr2−/− mice is rescued by adjuvants without improvement in germinal center development. Journal of immunology 165, 3119–3127.

Yuseff, M.I., Pierobon, P., Reversat, A., and Lennon-Dumenil, A.M. (2013). How B cells capture, process and present antigens: a crucial role for cell polarity. Nat Rev Immunol. 13, 475–486. doi: 410.1038/nri3469.

Zhu, Q., Zhang, M., Shi, M., Liu, Y., Zhao, Q., Wang, W., Zhang, G., Yang, L., Zhi, J., Zhang, L.*, et al.* (2016). Human B cells have an active phagocytic capability and undergo immune activation upon phagocytosis of Mycobacterium tuberculosis. Immunobiology. 221, 558–567. doi: 510.1016/j.imbio.2015.1012.1003. Epub 2015 Dec 1019.

Zimmerman, L.M., Vogel, L.A., Edwards, K.A., and Bowden, R.M. (2010). Phagocytic B cells in a reptile. Biol Lett. 6, 270–273. doi: 210.1098/rsbl.2009.0692. Epub 2009 Oct 1021.

