## Supplemental Information for "Recreation of an antigen-driven germinal center in vitro by providing B cells with phagocytic antigen"

### SUPPLEMENTAL FIGURE LEGENDS

**Suppl. Figure 1.** Phagocytosis of bead-bound antigen by B cells induces the expression of Tfh markers and release of cytokines by OT-2 CD4$^+$ T cells. **(A)** Non-transgenic B cells from WT and *Rhog*$^{-/-}$ mice were preincubated with 1 μm beads coated with IgM+OVA or IgM alone, at a 20:1 bead/cell ratio, and co-cultured for 3 days with OT-2 T cells (1:1 B/T cell ratio). Overlay histogram to the left show the upregulation of CD40 expression in gated WT B220$^+$ B cells compared to their RhoG-deficient counterparts. Bar plot to the right represents mean ± S.D. ($n$ = 3): * $p$<0.05; ** $p$<0.005 (unpaired Student's t test). **(B)** Differentiation of naïve B cells from B1-8$^{hi}$ transgenic WT and *Rhog*$^{-/-}$ mice to GC B cells was followed along 7 days of in vitro culture with beads coated with either NIP-OVA or NP-CGG and OT-2 T CD4$^+$ T cells. The percentage of GC B cells was calculated according to high GL7 and low CD95 expression (GL7$^+$CD95$^+$). Line plot below represents mean ± S.D. ($n$ = 3). **(C)** Proliferation of naïve OT-2 CD4$^+$ T cells in response to antigen presentation by B cells preincubated with the indicated doses of 1 μm and 3 μm beads coated with NIP-OVA. The dose of beads is normalized according to their exposed surface considering them as spheres. The bead/B cell ratios for 1 μm beads used were 0.1:1 to 3:1; for 3 μm beads were: 0.033:1 to 3.3:1. T cell proliferation was calculated by CTV dilution at day 4 after stimulation. Data represents mean ± S.D. ($n$ = 3). **(D)** Induction of TFH marker (PD1 and CXCR5; PD1 and ICOS) expression in OT-2 CD4$^+$ T cells after 4 days of culture with WT B cells as in (A). Data represents mean ± S.D. ($n$ = 3). ). **(E)** Cytokine release by OT-2 CD4$^+$ T cells incubated with either WT or RhoG-deficient B cells preincubated with a 3:1 1 μm bead/B cell ratio of beads coated with NIP-OVA, or without beads. Cell supernatants were collected at day 7 and cytokine content determined by ELISA. Data represents mean ± S.D. ($n$ = 3).

**Suppl. Figure 2.** Generation of somatic mutations in IgH V genes in conditions of GC formation in vitro. **(A)** Number of IgH nucleotide mutations and frequency of non-silent mutations in the IgH V sequence of sorted B cells from B1-8$^{hi}$ transgenic WT or RhoG-deficient mice stimulated with a 3:1 bead/cell ratio of 1 μm beads coated with NP-OVA or with 100 ng/ml of soluble NP-

OVA and co-cultured for 7 days with OT-2 T cells. **(B)** Sequences of the B1-8Vh genes with amino acid replacement mutations detected in the NIP-OVA bead stimulated cultures and their distribution referred to the three CDR regions.

**Suppl. Figure 3.** Generation of a GC reaction by phagocytic B cells and helper T cells does not require a third cell type. **(A)** Sorting of naïve follicular B cells from B1-8$^{hi}$ transgenic WT mice and of naïve CD4$^+$ T cells from OT-2 transgenic WT mice. The Dump channel contain the CD11b$^+$ and CD43$^+$ cells (for follicular B cell sorting) and the B220$^+$, CD11b$^+$, CD8$^+$, NK1.1$^+$, F480$^+$ cells (for CD4$^+$ T cells). **(B)** Sorted B and T cells as in (A) were incubated with either 1 μm beads coated with NIP-OVA or NP-CGG (3:1 bead/B cell ratio), or with 100 ng/ml soluble NIP-OVA for 4 days at a 1:1 B/T cell ratio. FACS contour-plots to the left show the appearance of a double positive (CD95$^+$ GL7$^+$) population in gated B220$^+$ B cells. Bar plot to the right represents mean ± S.D. ($n$ = 3): * $p$<0.05; ** $p$<0.005 (unpaired Student's t test). **(C)** Detection of high-affinity anti-NP Igs in supernatants from B and T cell cultures prepared as in (A) and (B) incubated for 7 days (3:1 1 μm bead/B cell ratio) and soluble antigen concentration. Data represent the mean ± S.D. ($n$ = 3).

### 2.-SUPPLEMENTARY EXPERIMENTAL PROCEDURES

#### KEY RESOURCES TABLE

| REAGENT or RESOURCE | SOURCE | IDENTIFIER |
|---|---|---|
| Antibodies | | |
| Anti-Actin | | |
| Anti-AKT (clone 9272) | Cell Signaling | Cat#9272 |
| Anti-CD45R APC (clone RA3-6B2) | BD Pharmigen | Cat#553092 |
| Anti-CD45R Biotin (clone RA3-6B2) | BD Pharmigen | Cat#553085 |
| Anti-CD45R FITC (clone RA3-6B2) | BD Pharmigen | Cat#553088 |
| Anti-CD45R V450 (clone RA3-6B2) | BD Pharmigen | Cat#560473 |
| Anti-Bcl6 647 (clone K112-91) | BD Pharmigen | Cat#561525 |
| Anti-Bcl6 PE (clone K112-91) | BD Pharmigen | Cat#561522 |
| Anti-Blimp1 PE (clone 5E7) | BD Pharmigen | Cat#564268 |
| Anti-CD4 647 (clone RM4-5) | BD Pharmigen | Cat#557681 |
| Anti-CD4 PerCP (clone RM4-5) | BD Pharmigen | Cat#553052 |
| Anti-CD8 Biotin (clone 53-6.7) | BD Pharmigen | Cat#553029 |

| | | |
|---|---|---|
| Anti-CD11c Biotin (clone HL3) | BD Pharmigen | Cat#553800 |
| Anti-CD16/32 purified (clone 2.4G2) | BD Pharmigen | Cat#553141 |
| Anti-CD19 PE (clone 1D3) | eBiosciences | Cat#12-0193 |
| Anti-CD19 PE-Cy7 (clone 1D3) | BD Pharmigen | Cat#561739 |
| Anti-CD25 APC (clone 3C7) | BD Pharmigen | Cat#557192 |
| Anti-CD40 PE-Cy5 (clone 3/23) | Biolegend | Cat#124617 |
| Anti-CD43 Biotin (clone S7) | BD Pharmigen | Cat#553269 |
| Anti-CD45.1 APC-Cy7 (clone A20) | BD Pharmigen | Cat#560579 |
| Anti-CD45.2 APC (clone 104) | BD Pharmigen | Cat#558702 |
| Anti-CD86 PE-Cy5 (clone GL1) | eBiosciences | Cat#15-0862 |
| Anti-CD95 FITC (clone Jo2) | BD Pharmigen | Cat#554257 |
| Anti-CD95 PE-Cy7 (clone Jo2) | BD Pharmigen | Cat#557653 |
| Anti-CD138 APC (clone 281-2) | BD Pharmigen | Cat#558626 |
| Anti-CD154 PE (clone MR1) | BD Pharmigen | Cat#561719 |
| Anti-CD275 Biotin (clone HK5.3) | eBiosciences | Cat#13-5985 |
| Anti-CD278 PE (clone 7E.17G9) | Biolegend | Cat#117405 |
| Anti-CD279 (PD1) (clone J43) | eBiosciences | Cat#11-9985-85 |
| Anti-CXCR4 Biotin (clone 2B11) | eBiosciences | Cat#13-9991-80 |
| Anti-CXCR5 Biotin (clone 2G8) | BD Pharmigen | Cat#551960 |
| Anti-F4/80 Biotin (clone BM8) | Biolegend | Cat#123105 |
| Anti-Gr1 Biotin (clone RB6-8C5) | BD Pharmigen | Cat#553125 |
| Anti-GL7 647 (clone GL7) | BD Pharmigen | Cat#561529 |
| Anti-GL7 FITC (clone GL7) | BD Pharmigen | Cat#553666 |
| Anti-IgD Biotin (clone 11-26c) | eBiosciences | Cat#13-5993-81 |
| Anti-IgD FITC (clone 11.26c) | BD Horizon | Cat#562022 |
| Anti-IgD V450 (clone 11.26c) | BD Horizon | Cat#560869 |
| Anti-goat IgGs FITC | Jackson Immunoresearch | Cat#705-546-147 |
| Anti-kappa Biotin (clone RMK-12) | Biolegend | Cat#407204 |
| Anti-IgM PE (clone II/41) | eBiosciences | Cat#12-5790-81 |
| Anti-IgM APC (clone II/41) | eBiosciences | Cat#17-5790-82 |
| Anti-IgM F(ab')2 | Jackson Immunoresearch | Cat#115-006-075 |
| Anti-NK1.1 Biotin | BD Pharmigen | Cat#553163 |
| Anti-pAkt (S473) (clone D9E) | Cell Signaling | Cat#4060 |
| Anti-pCD79$\alpha$ (Y182) | Cell Signaling | Cat#5173 |
| Anti-pErk (T202/Y204) | Cell Signaling | Cat#9101 |
| Anti-pS6 (S240/244) (cloneD68F8) | Cell Signaling | Cat#5364) |
| Anti-pSYK (Y525/Y526) | Cell Signaling | Cat#2711 |
| Anti-V$\alpha$2 PercP/Cy5.5 (clone B20.1) | Biolegend | Cat#127813 |
| Biological Samples | | |
| Chemicals, Peptides, and Recombinant Proteins | | |
| Polybead Carboxylate 1 $\mu$m | Polysciences | Cat#08226-15 |
| Polybead Carboxylate 3 $\mu$m | Polysciences | Cat#09850 |
| FluoSpheres Carboxylate 1 $\mu$m Crimson (625/645) | ThermoFischer | Cat#F8816 |
| Fluoresbrite Carboxylate 3 $\mu$m Y/O (529/546) | Polysciences | Cat#19393 |
| Fluoresbrite Carboxylate 1 $\mu$m Y/G (441/486) | Polysciences | Cat#15702 |

| | | |
|---|---|---|
| Fluoresbrite Carboxylate 3 μm Y/G (441/486) | Polysciences | Cat#17147 |
| Fluoresbrite Carboxylate 10 μm Y/G (441/486) | Polysciences | Cat#18142 |
| Cell Trace Violet | Life Technology | Cat#C34557 |
| DAPI | Merck | Cat#268298 |
| Ovalbumin | SIGMA | Cat#A5503 |
| NIP(15)-Fluorescein-BSA | Biosearch Technology | Cat#N-5040F-10 |
| NIP(7)-BSA | Biosearch Technology | Cat#N-5050L-10 |
| NIP(41)-BSA | Biosearch Technology | Cat#N-5050H-10 |
| NP(25)-CGG | Biosearch Technology | Cat#N-5055C-5 |
| NP(36)-PE | Biosearch Technology | Cat#N-5070-1 |
| Phalloïdin-TRICT | Sigma | Cat#P-1951 |
| Phalloïdin-647 | ThermoFisher | Cat#A-22287 |
| Streptavidin-PercP | BD Pharmigen | Cat#554064 |
| Streptavidin-APC | BD Pharmigen | Cat#554067 |
| Streptavidin-APC-Cy7 | BD Pharmigen | Cat#554063 |
| HIV-1 p17/p24/gp120 | Jena Biosciences | Cat#PR-1202 |
| Imject Alum | Thermo Scientific | Cat#77161 |
| Ampicillin | Sigma | Cat#A1593 |
| IL-4 | Peprotech | Cat#214-14 |
| IL-21 | Peprotech | Cat#210-21 |
| Sheep Red Blood Cells (SRBC) | Oxoid | Cat#SR0051 |
| Critical Commercial Assays | | |
| SBA Clonotyping system-HRP | Southern Biotech | Cat#5300-04 |
| RNeasy Plus Mini Kit | QIAGEN | Cat#74136 |
| GoTaq qPCR Master Mix | PROMEGA | Cat#A-6002 |
| QIAamp DNA kit | QIAGEN | Cat#51304 |
| Foxp3 / Transcription Factor Staining Buffer Set | eBiosciences | Cat#00-5523-00 |
| Mouse IL-2 Flex Set | BD Biosciences | Cat#558297 |
| MS/Rat Soluble Protein master Buffer Kit | BD Biosciences | Cat#558266 |
| Mouse IL-21 Flex Set | BD Biosciences | Cat#560160 |
| IL-10 CBA Flex Set | BD Biosciences | Cat#558300 |
| IL-6 CBA Flex Set | BD Biosciences | Cat#558301 |
| IL-4 CBA Flex Set | BD Biosciences | Cat#558298 |
| JetPei DNA transfection | Polyplus | Cat#101-01N |
| Deposited Data | | |
| Raw data files for DNA sequences | NCBI Gene Expression Omnibus | |
| Experimental Models: Cell Lines | | |
| HEK293T | ATCC | Cat#CRL-3216 |
| MOLT-4 | ATCC | Cat#CRL-1582 |
| Mouse: primary B lymphocytes | This paper | N/A |
| Mouse: primary T lymphocytes | This paper | N/A |
| Experimental Models: Organisms/Strains | | |
| E.Coli DH5α | CBMSO fermentation core facility | N/A |

| | | |
|---|---|---|
| Mouse: C57BL/6 | Envigo | |
| Mouse: RRas2-/- | (Delgado et al., 2009) | N/A |
| Mouse: RhoG-/- | (Vigorito et al., 2004) | N/A |
| Mouse: B6.129S7-*Rag1*$^{tm1Mom}$/J (Rag-/-) | Provided by Cesar Cobaleda | Stock number: 002216 |
| Mouse: B6.Cg-Tg(TcraTcrb)425Cbn/J (OT-II) | Provided by Carlos Ardavín | Stock number: 004194 |
| Mouse: B6.SJL-*Ptprc*$^{a}$ *Pepc*$^{b}$/BoyJ (CD45.1) | Provided by Carlos Ardavín | Stock number: 002014 |
| Recombinant DNA | | |
| pHRSin-GFP | Provided by J. A. Pintor | (Demaison et al., 2002) |
| pCMV-NL4.3 | This study | N/A |
| psPAX2 | This study | Addgene plasmid: 12260 |
| pMD2.G | This study | Addgene plasmid: 12259 |
| Sequence-Based Reagents | | |
| **Bcl6** FW GGAAGTTCATCAAGGCCAGT / RV GACCTCGGTAGGCCATGA | This paper | N/A |
| **Bcl2** FW GTACCTGAACCGGCATCTG / RV GGGGCCATATAGTTCCACAA | This paper | N/A |
| **Blimp1** FW GGCTCCACTACCCTTATCCTG / RV TTTGGGTTGCTTTCCGTTT | This paper | N/A |
| **B1-8Vh** FW CCATGGGATGGAGCTGTATCATCC / RV GAGGAGACTGTGAGAGTGGTGCC | (Shih et al. 2002) | N/A |
| **HPRT** FW / TCCTCCTCAGCAAGCTTTT / RV CCTGGTTCATCATCGCTAATC | This paper | N/A |
| **GAPDH** FW CTCCCACTCTTCCACCTTCG / RV CATACCAGGAAATGAGCTTGACAA | This paper | N/A |
| Software and Algorithms | | |
| FloWJo | FloWJo | N/A |
| Fiji (ImageJ) | https://fiji.sc/ | N/A |
| FACS Diva software | BD Biosciences | N/A |
| Adobe Photoshop and Illustrator CS4 | Adobe | N/A |
| Other | | |

### CONTACT FOR REAGENT AND RESOURCE SHEARING

### METHOD DETAIL

#### Cell preparation and purification

The lymph nodes and spleen from 6-8 weeks mice were homogenized with 40 μm strainers and washed in phosphate-buffered saline (PBS) containing 2% (vol/vol) fetal bovine serum (FBS). Spleen cells were resuspended for 3 minutes in AcK buffer (0.15 M $NH_4Cl$, 10mM $KHCO_3$, 0.1 mM EDTA, pH 7.2-7.4) lo lyse the erythrocytes and washed in PBS 2% FBS. B cells from spleen were negatively selected using a combination of biotinylated anti-CD43 and anti-CD11b antibodies and incubation with streptavidin beads (Dynabeads Invitrogen) for 30 minutes and separated using Dynal Invitrogen Beads Separator. B1-8$^{hi}$ B cells were purified using biotinylated anti-CD43, anti-CD11b and anti-kappa antibodies. OT-2 T cells from lymph nodes and spleen were purified using a mix of biotinylated antibodies: anti-B220, anti-CD8, anti-NK1.1, anti-CD11b, anti-GR1, and anti-F4/80. Splenic and lymph node B and T cells were maintained in RPMI 10% FBS supplemented with 2 mM L-glutamine, 100U/ml penicillin, 100 U/ml streptomycin, 20 μm β-mercaptoethanol and 10 mM sodium pyruvate.

#### Real-time PCR

A total of $5x10^6$ purified B1-8$^{hi}$ WT or *Rhog*$^{-/-}$ B2 cells were cultured with purified OT2 (ratio 1:1) and different BCR stimuli (NIP-OVA bound to 1 μm beads, 100 ng/mL soluble NIP-OVA or NP-CGG bound to 1 μm beads) in a 6 well flat-bottom plate. After 7 days, B cells were sorted (FACSAria Fusion (BSC II)) and their RNA was isolated using the RNAeasy Plus Mini Kit (QIAGEN). cDNA was synthetized with SuperScript III (Invitrogen) using Oligo-dT primers. Quantitative real-time PCR was performed in triplicate using the reverse transcription reaction with SYBR Green PCR Master Mix, gene-specific primers and ABI 7300 Real Time PCR System. Obtained cycle threshold (Ct) values were used to calculate mRNA levels relative to the HPRT and 18S RNA expression using the 2-ΔΔCt method.

#### BCR downmodulation

Purified B1-8$^{hi}$ B cells were resuspended in RPMI plus 10% FBS at a density of 2.5 x$10^5$ cells /well in a p96 V-bottom plate in a total volume of 50 μL. Stimulus (NIP-OVA bound to beads in a 3:1 ratio or soluble NIP-OVA at 100 ng/ml concentration) were added to the wells and incubated at 37ºC for different time points. After appropriate incubation time, cells were stained for IgM, CD19 and B220 at 0ºC and analysed by FACS.

**Confocal microscopy**

For immunofluorescence analysis of B:T cell clusters, purified B cells were cultured with purified OT2 T cells for 4 or 7 days in 6 well flat-bottom plates with either 1μm beads coated with anti-IgM plus ovalbumin or NIP-OVA or 100 ng/ml soluble NIP-OVA. Afterwards, cells were fixed in 4% paraformaldehyde for 20 minutes and transferred to poly-L-lysine treated coverslips. Cells were stained for 1 hour at 0ºC with antibodies specific for B220, CD4 and GL7. For analysis of intracellular phosphorylation, purified B1-8$^{hi}$ B cells were starved for one hour in RPMI plus 20mM HEPES-HCl pH=7.4, and subsequently stimulated with NIP-OVA-coated fluorescent-beads (3 bead:B cell ratio) or soluble NIP-OVA (100ng/mL). Cells were fixed in 4% paraformaldehyde at 0ºC for 20 minutes to stop the stimulus, washed with Tris-buffered saline (TBS), and adhered to poly-L-lysine treated coverslips. Extracellular staining for B220 was performed in TBS for 1 hour at 0 ºC. After that, cells were stained for anti-phospho-Syk and anti-phospho-Igα as suggested by the manufacter (Cell Signaling). Confocal images were acquired with a Zeiss LSM710 system and a Zeiss AxioObserver LSM710 Confocal microscopes.

**Measurement of actin polymerization**

Purified B1-8$^{hi}$ B cells were starved as above and stimulated with either NIP-OVA bound to beads or soluble NIP-OVA for different times at 37 ºC. After that, cells were washed once in PBS plus 1 % BSA at 37 ºC and fixed with 4% paraformaldehyde for 10 minutes at room temperature. Extracellular staining for B220 was performed in PBS plus 1% BSA. After washing, cells were permeabilized in 4% paraformaldehyde plus 0.1% Triton X-100 for two minutes at room temperature. Phalloïdin-Alexa488 was diluted 1:200 in PBS plus 1%BSA and incubated with cells for 1 hour. After washing, cells were analysed by Flow Cytometry (FACS Canto II).

**NP saturation assay**

A total of $1x10^5$ purified B1-$8^{hi}$ B cells/well were plated in a V bottom 96 well plates and incubated with different doses of stimuli for one hour at 0ºC. Cells were washed once with cold PBS plus 1% BSA and stained with NP(36)-PE at 2.5 μg/ml in 50 μL per well at 0ºC for 30 minutes. Cells were washed once with cold PBS plus 1% BSA and analyzed by Flow Cytometry (FACS Canto II).

**Immunoblot analysis of B cell activation**

Purified B1-$8^{hi}$ B cells were resuspended in RPMI plus 20 mM Hepes and left in starving conditions for 1 hour. Cells were stimulated at different time-points with NIP-OVA bound-beads (ratio 3:1 beads/B cell) or 100ng/mL soluble NIP-OVA. After stimulation, cells were lysed in Brij96 lysis buffer containing protease and phosphatase inhibitors (1% Brij96, 140 mM NaCl, 10 mM Tris–HCl [pH 7.8], 10 mM iodoacetamide, 1 mM PMSF, 1 μg/ml leupeptin, 1 μg/ml aprotinin, 1 mM sodium orthovanadate, 20 mM sodium fluoride and 5 mM of MgCl2). Immunoblotting was performed as described previously (Martinez-Martin et al., 2009)

**Somatic hypermutation**

Purified WT or RhoG-/- B1-8 B cells were cultured with purified OT-II T cells (ratio 1:1) and soluble NIP-OVA (100ng/mL) or bead-bound NIP-OVA (ratio 3:1 beads/Bcell). After 7 days of culture, B1-8 cells were sorted and their genomic DNA was extracted using QIAamp DNA kit (QIAGEN). B1-8Vh genes were amplified by PCR with the Expand High Fidelity (Roche) and the primers forward 5'-CCATGGGATGGAGCTGTATCATCC-3' and reverse 5'-GAGGAGACTGTGAGAGTGGTGCC-3 as described previously (Shih et al. 2002). PCR products were subcloned into PCR2.1 vector (Invitrogen). DH5α bacteria were transformed with the subcloned products. Individual DH5α clones grown in LB+Ampicillin were selected for sequencing using the SUPREMERUN 96 system of GATC.

**Interleukin measurement**

Purified WT B1-8$^{hi}$ B cells were cultured together with OT2 T cells and two different bead sizes (1 μm and 3 μm) bound to NIP-OVA at different doses. After 4 days, the supernatant was used to measure cytokines with a BD Cytometric Bead Array (CBA) Kit as indicated by the manufacturer. In cultures of WT or RhoG-deficient B1-8$^{hi}$ B cells with purified OT2 T cells and NIP-OVA 1 μm bead-bound, the supernatant was obtained after 7 days.

### References

Delgado, P., Cubelos, B., Calleja, E., Martinez-Martin, N., Cipres, A., Merida, I., Bellas, C., Bustelo, X.R., and Alarcon, B. (2009). Essential function for the GTPase TC21 in homeostatic antigen receptor signaling. Nature immunology *10*, 880-888.
Demaison, C., Parsley, K., Brouns, G., Scherr, M., Battmer, K., Kinnon, C., Grez, M., and Thrasher, A.J. (2002). High-level transduction and gene expression in hematopoietic repopulating cells using a human immunodeficiency [correction of imunodeficiency] virus type 1-based lentiviral vector containing an internal spleen focus forming virus promoter. Hum Gene Ther *13*, 803-813.
Vigorito, E., Bell, S., Hebeis, B.J., Reynolds, H., McAdam, S., Emson, P.C., McKenzie, A., and Turner, M. (2004). Immunological function in mice lacking the Rac-related GTPase RhoG. Molecular and cellular biology *24*, 719-729.
