## Supplementary figures and images for "Recreation of an antigen-driven germinal center in vitro by providing B cells with phagocytic antigen"

### Supplemental Figure 1

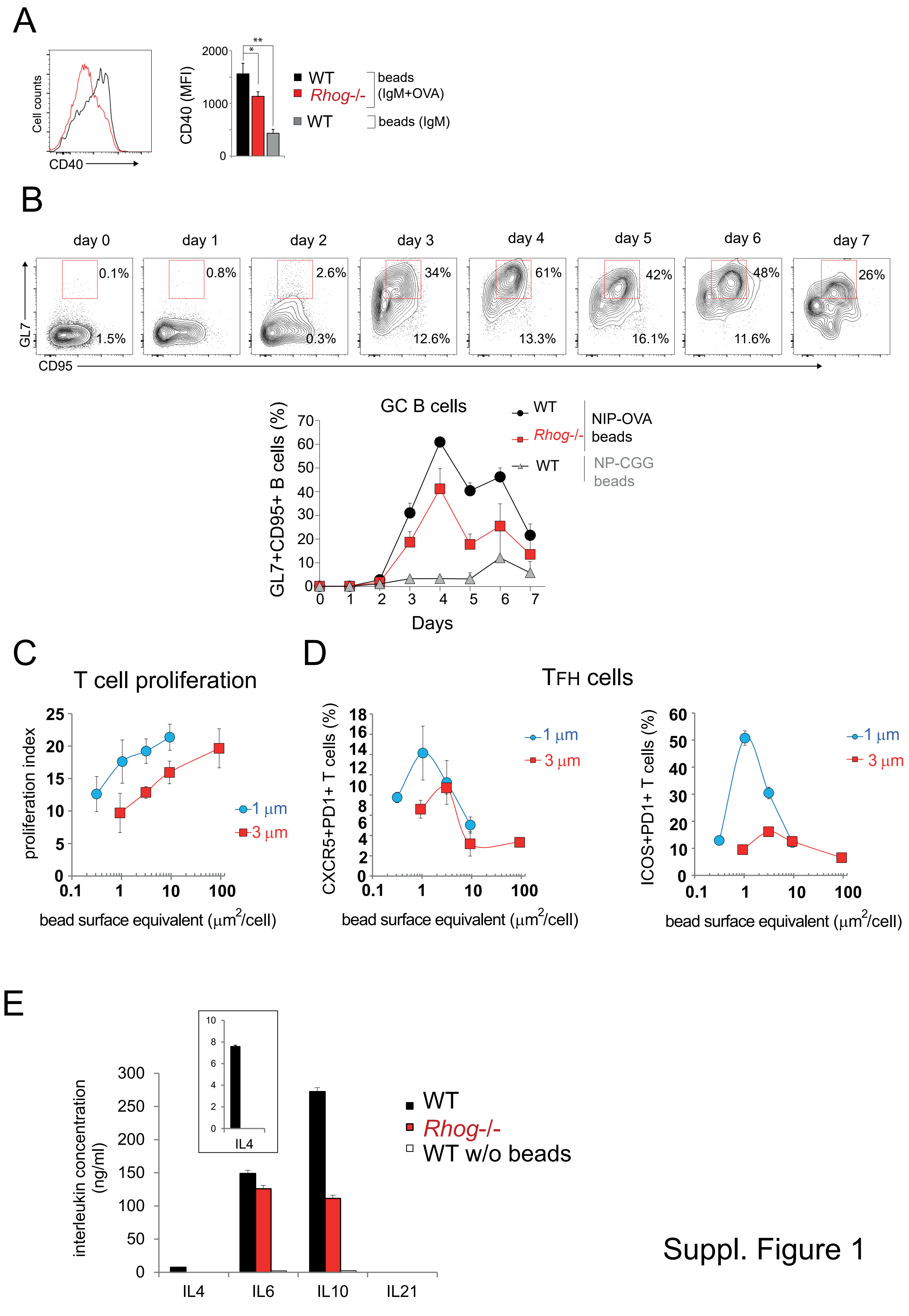


Suppl. Figure 1

### Supplemental Figure 3

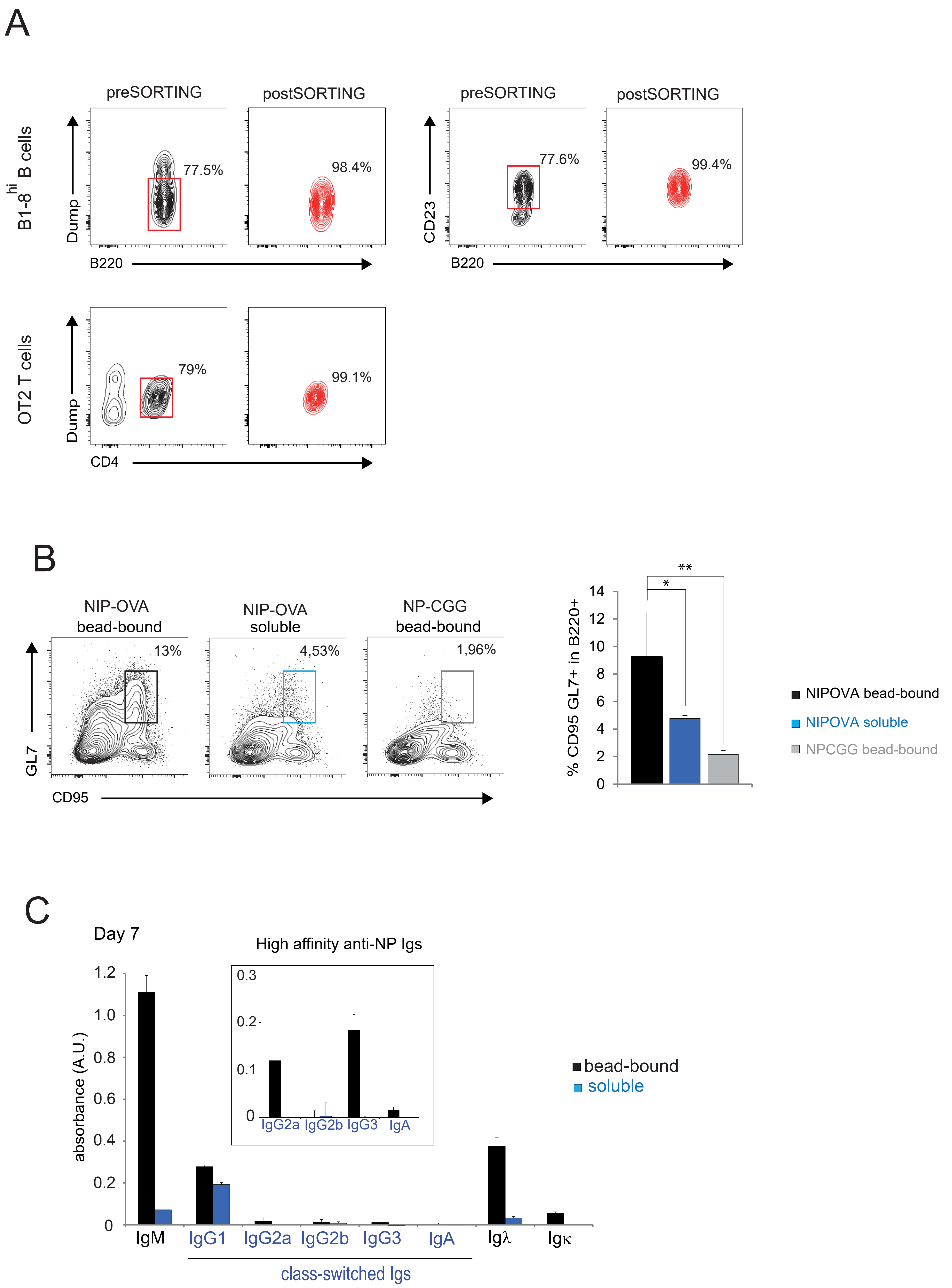
