## Supplemental Figure 2 for "Recreation of an antigen-driven germinal center in vitro by providing B cells with phagocytic antigen"

## A

| | no. sequences | no. bases | no. mutations | mutation rate (nucleotide) x10E-5 | mutation rate (protein) |
|---|---|---|---|---|---|
| non-stim WT | 116 | 50209 | 2 | 4.0 | 0.000000006 |
| beads WT | 360 | 154607 | 49 | 31.7 | 0.0006 |
| soluble WT | 232 | 98725 | 21 | 21.3 | 0.0004 |
| beads RhoG KO | 47 | 20194 | 1 | 4.9 | 0.0 |

## B

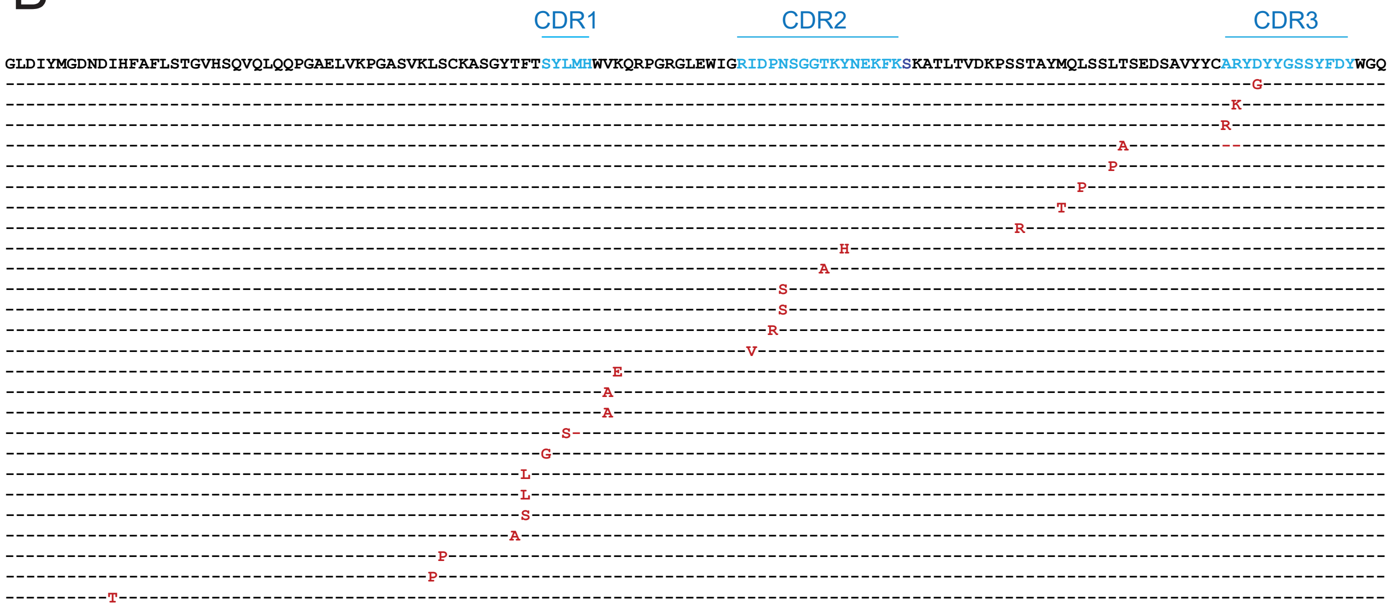


Suppl. Fig. 2
